# MmcA is an electron conduit that facilitates both intracellular and extracellular electron transport in *Methanosarcina acetivorans*

**DOI:** 10.1101/2023.04.20.537704

**Authors:** Dinesh Gupta, Keying Chen, Sean J. Elliott, Dipti D. Nayak

## Abstract

Methanogens are a diverse group of Archaea that couple energy conservation to the production of methane gas. While most methanogens have no alternate mode of energy conservation, strains like *Methanosarcina acetivorans* are known to also conserve energy by dissimilatory metal reduction (DSMR) in the presence of soluble ferric iron or iron-containing minerals. The ecological ramifications of energy conservation decoupled from methane production in methanogens are substantial, yet the molecular details are poorly understood. In this work, we conducted *in vitro* and *in vivo* studies with a multiheme *c*-type cytochrome (MHC), called MmcA, to establish its role during methanogenesis and DSMR in *M. acetivorans*. MmcA purified from *M. acetivorans* can donate electrons to methanophenazine, a membrane-bound electron carrier, to facilitate methanogenesis. In addition, MmcA can also reduce Fe(III) and the humic acid analog anthraquinone-2,6-disulfonate (AQDS) during DSMR. Furthermore, mutants lacking *mmcA* have slower Fe(III) reduction rates. The redox reactivities of MmcA are consistent with the electrochemical data where MmcA displays reversible redox features ranging from -100 to -450 mV versus SHE. MmcA is prevalent in members of the Order *Methanosarcinales* but does not belong to a known family of MHCs linked to extracellular electron transfer, bioinformatically, and instead forms a distinct clade that is closely related to octaheme tetrathionate reductases. Taken together, this study shows that MmcA is widespread in methanogens with cytochromes where it acts as an electron conduit to support a variety of energy conservation strategies that extend beyond methanogenesis.

## Significance Statement

Methane is likely to contribute more to global warming in the next decade than carbon dioxide. The primary source of biogenic methane is methanogens, a group of Archaea that conserve energy via methanogenesis. Here, we show that methanogens like *Methanosarcina acetivorans* can conserve energy by metal respiration instead of methanogenesis using a multiheme *c*-type cytochrome called MmcA. In addition to its role in intracellular electron transport during methanogenesis, MmcA can also transfer electrons to extracellular electron acceptors like Fe(III), which enables *M. acetivorans* to proliferate without producing methane. Therefore, the presence of MmcA has a remarkable impact on the physiology of methanogens with cascading ramifications on global methane emissions.

## Introduction

Methanogens are a polyphyletic group of Archaea that couple energy conservation to the production of methane gas (1, 2). A consensus view based on the hundreds of isolates studied so far is that they are all obligate methanogens (3). This observation is supported by the argument that methanogenesis requires a large number of unique cofactors, enzymes and bioenergetic complexes that may not be compatible with other modes of energy conservation in the cell (2–5). That said, there are a few studies that allude to the possibility that methanogens might be able to conserve energy without producing methane, especially in the presence of extracellular electron acceptors like humic acids or iron containing minerals (6–13). Under these circumstances, methanogens are hypothesized to conserve energy through dissimilatory metal reduction (DSMR) i.e., by coupling the oxidation of inorganic or organic electron donors with the aforementioned electron acceptors (12, 13). The phylogenetic breadth of methanogens that are capable of DSMR is unclear. While some studies suggest that this trait is inherent to all methanogens, others indicate that it is restricted to methanogens with cytochromes (and an electron transport chain) such as members of the Order *Methanosarcinales* (6–13). Moreover, some studies show that the presence of extracellular electron acceptors completely inhibits methanogenesis whereas others show that DSMR and methanogenesis can happen simultaneously (13–18). Altogether, it seems likely that some methanogens encode alternate energy conservation strategies, but the molecular details of these pathways need to be uncovered in order to glean their taxonomic breath and ecological significance.

Some recent work indicates that *c-*type cytochromes (referred to as cyt(s) *c* throughout the text) might facilitate DSMR through extracellular electron transfer (EET) in the model methanogen, *Methanosarcina acetivorans* (12, 19, 20). Most notably, a mutant of *M. acetivorans* lacking a multiheme cyt *c* (MHC) called MmcA was unable to grow in media with the humic acid analog anthraquinone-2,6-disulfonate (AQDS) as the sole electron acceptor in the presence of 2-bromoethanesulfonic acid (BES), a potent inhibitor of methanogenesis (12). While this initial study showed that MmcA is not required during methanogenic growth, follow up work provides compelling evidence that MmcA is also critical for energy conservation during methanogenesis (21, 22), which aligns with its proposed role in the <u>R</u>hodobacter <u>N</u>itrogen <u>F</u>ixation (Rnf) complex (23–25). In methanogens, the Rnf complex couples the energetically favorable electron transfer from reduced ferredoxin to the membrane-bound electron carrier methanophenazine (MP) to the translocation of Na^+^ ions across the cytoplasmic membrane (Supplementary Figure 1) (23–25). Mutants lacking *mmcA* are also devoid of a functional Rnf complex and are completely incapable of methanogenic growth on substrates like acetate (21, 22). Altogether, while it is clear that MmcA is an integral component of the electron transport chain (ETC) in *M. acetivorans*, there is lingering ambiguity about its role in supporting different energy conservation strategies (Supplementary Figure 2).

In this study we use a combination of *in vitro* and in *vivo* techniques to delineate two distinct biochemical role(s) for MmcA that depend on the environmental context. First, we provide conclusive evidence that MmcA is a membrane-associated heptaheme cyt *c.* Next, using MmcA purified from *M. acetivorans,* we demonstrate that this protein acts as an MP-reductase to facilitate intracellular electron transport during methanogenesis. In addition, we show MmcA can also transfer electrons to AQDS and ferric iron. Thus, if these electron acceptors are present in the medium, methanogens like *M. acetivorans* can perform DSMR. Finally, we also show that the redox properties and evolutionary origins of MmcA are consistent with its role as low potential electron conduit that supports a wide range of energy conservation strategies in methanogens.

## Results

### Production and purification of MmcA from *M. acetivorans*

*MmcA* (locus tag: MA0658 or MA_RS03460) is the first gene of the Rnf operon (MA0658-MA0665 or MA_RS03460-MA_RS03495), and it encodes a 500-amino acid (aa) apo-protein with an N-terminal Sec signal peptide (1-24 aa) (Supplementary Figure 3). MmcA contains five canonical (CXXCH) and two non-canonical (CXXXCH and CXXXXCH) heme-binding motifs and the holo-protein is predicted to be a 476 aa-long heptaheme cyt *c* after processing the signal sequence (Supplementary Figure 3). MmcA is an essential component of the Rnf complex in *M. acetivorans* and is primarily found in the membrane fraction (Supplementary Figure 4) (23). Since MmcA lacks an identifiable transmembrane domain, we anticipate that it is tethered in the membrane through interactions with membrane integral proteins of the Rnf complex (26). To overexpress and purify MmcA from *M. acetivorans*, we generated an overexpression vector encoding a C-terminal tagged (3×FLAG tag and a twin-Strep tag) *mmcA* placed under the control of tetracycline-inducible promoter (21). Our previous work shows that the tag does not interfere with maturation of holo-MmcA and that the tagged protein can also rescue the growth defect of Δ*mmcA* mutant during growth on methylated compounds (21, 22). Multiple attempts to purify the C-terminal tagged MmcA using a Streptactin resin using either phosphate or Tris buffers (pH = 7.0-8.0) were unsuccessful (Supplementary Figure 5) (27, 28). However, by switching to an anti-Flag resin we were able to obtain ∼ 250 µg of protein per liter of culture (Supplementary Figure 6). We performed a peroxidase-based heme stain and an immunoblot with anti-Flag antibody to confirm that the purified protein contains covalently bound heme groups i.e. is a holo-form of MmcA (Fig. 1A). Purified MmcA has a distinct red color which is a hallmark of cyt *c* (Fig. 1A) and also has spectral features typical of cyt *c* with an absorption maximum at 408 nm (γ) and 530 nm in the oxidized state (in black) and 419.5 nm (γ), 523 nm (β), and 552 nm (α) in the reduced state (in red) (Fig. 1B). We also confirmed the presence of covalently attached heme groups in MmcA by performing a pyridine hemochrome assay and observed a characteristic 550-nm α peak (Fig. 1B, inset and Supplementary Figure 7). Taken together, these data show that purified MmcA from *M. acetivorans* is a heme-attached cyt *c* protein.

**Figure 1:**
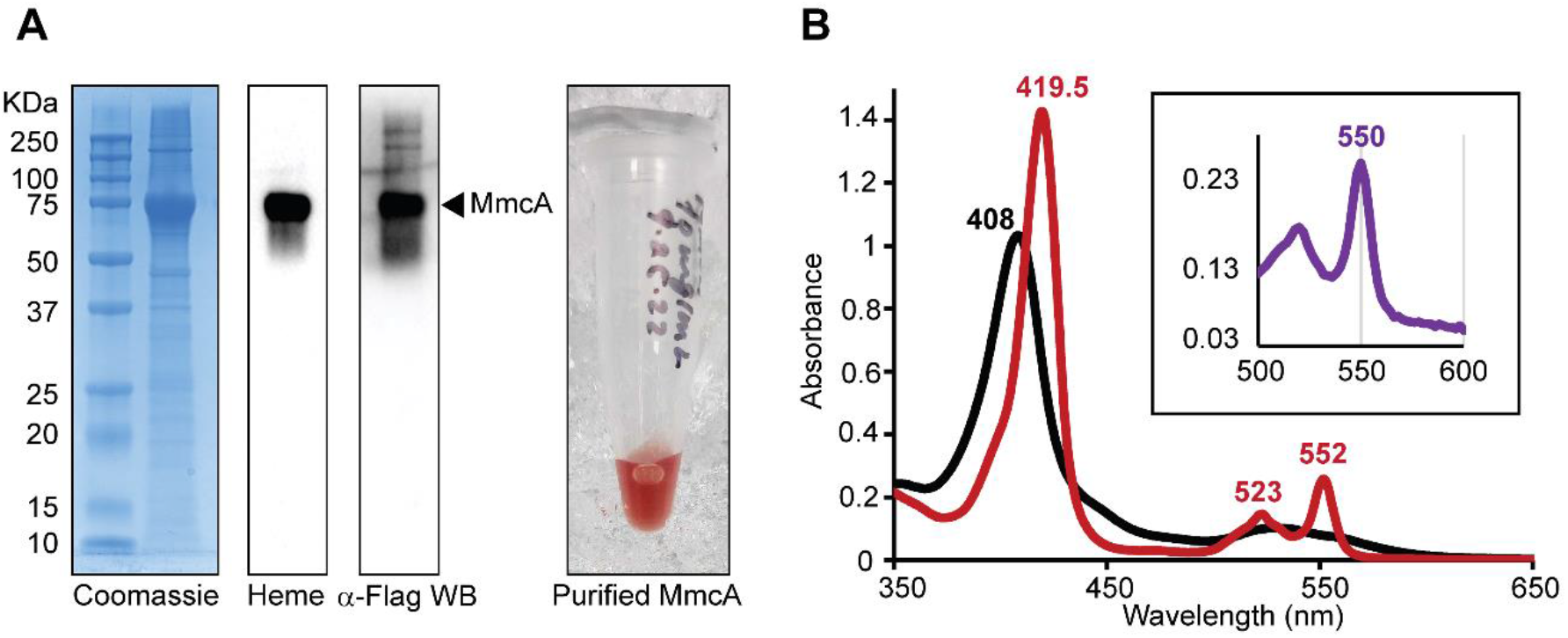
Purification and UV-vis characterization of MmcA. (**A**) Coomassie, heme staining and Western blot (WB) with anti (α)-Flag antibody of C-terminal 3×FLAG tagged MmcA from *Methanosarcina acetivorans*. MmcA purified under aerobic conditions and concentrated to *ca.* 100 µM has a reddish-brown color (**B**) UV-vis spectral analysis of purified MmcA in the oxidized state (black) and reduced with sodium dithionite (red). The inset shows the pyridine hemochrome assay of reduced MmcA with a characteristic alpha peak for *c*-type cytochromes at 550 nm. (Complete UV-vis spectra of the hemochrome assay is shown in Supplementary Figure 7).

### MmcA is a heptaheme cytochrome *c*

Although MmcA is predicted to be a heptaheme cyt *c*, the heme occupancy of this protein has not been experimentally validated. To this end, we performed LC-MS/MS analysis of chymotrypsin digested fragments of MmcA and observed that the seven peptides with the putative heme-binding motifs had an increase in mass of 615.17 Da corresponding to heme attachment (Table 1) (29). This observation confirms that MmcA is a heptaheme cyt *c*, and also demonstrates that the cyt *c* maturation (CCM) machinery of *M. acetivorans* can covalently attach heme to non-canonical heme binding motifs in MmcA (Supplementary Figure 3) (21). We did not detect the predicted signal peptide in either trypsin or chymotrypsin digested fragments of MmcA as it gets cleaved during the translocation of MmcA to the pseudo-periplasm of *M. acetivorans* (Supplementary Figure 8).

**Table 1:**
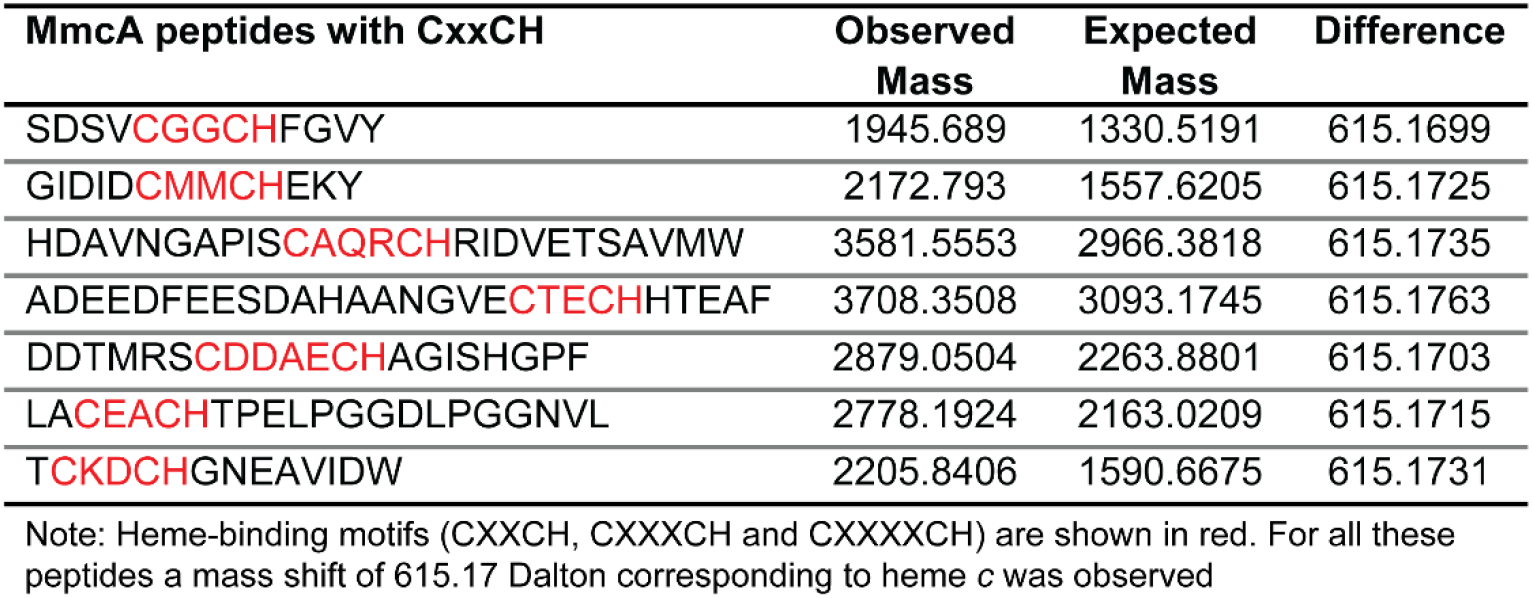
Chymotrypsin-digested MmcA-peptides with heme-binding motifs.

### MmcA is a methanophenazine reductase

While it has been hypothesized that MmcA mediates the final step in transferring electrons from the Rnf complex to membrane-bound electron-carrier MP (23, 24), experimental validation is still pending. To this end, we conducted spectroscopic analyses with purified MmcA to measure its MP-reductase activity (Fig. 2A). We added an excess of 2-hydroxyphenazine (2HP) (200 µM), a well-established soluble analog of MP (24, 30–33), to MmcA reduced with sodium dithionite under anaerobic conditions (Fig. 2B). The addition of 2HP oxidized MmcA as observed by a shift in its Soret peak from 419.5 nm to 408 nm and the disappearance of the α and β peaks of reduced-MmcA (Figure 2B; Supplementary Figure 8). These *in vitro* data provide conclusive evidence that MmcA plays the role of an MP reductase in the Rnf complex from *M. acetivorans*.

**Figure 2:**
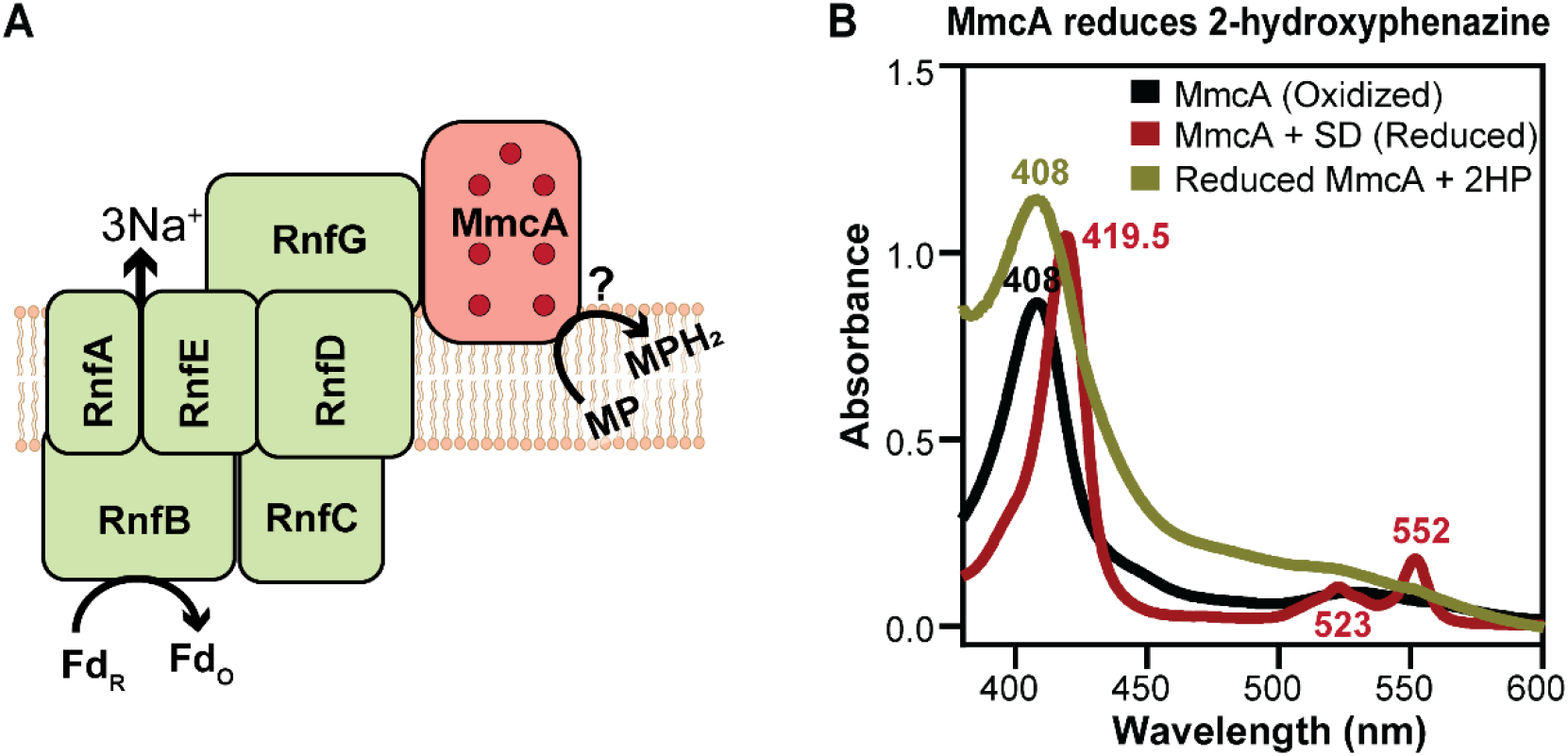
MmcA can function as a methanophenazine (MP) reductase. **(A)** Schematic of the MmcA-Rnf complex where the multiheme cytochrome MmcA mediates the final step in the transfer of electrons from the complex to the membrane-bound electron carrier methanophenazine (MP). (**B**) Spectral analysis of MmcA-mediated reduction of the soluble MP analog, 2-hydroxyphenazine (2HP) under anaerobic conditions. Addition of 2HP to MmcA reduced with sodium dithionite (SD; red) leads to the oxidation of MmcA (olive green) as observed by the appearance of a characteristic Soret peak at 408 nm indicative of the oxidized protein (black).

### MmcA can donate electrons to extracellular electron acceptors like ferric iron and humic acid analogs

Previous studies have shown that a mutant of *M. acetivorans* lacking MmcA fails to reduce AQDS (12), but these genetic data do not imply that MmcA interacts with and reduces AQDS and other electron acceptors (Fig. 3A). We tested if MmcA can reduce AQDS or ferric iron [either ferric chloride (FeCl_3_) or ferricyanide (K_3_[Fe(CN)_6_])] by monitoring the redox state of MmcA after adding an excess of each of these electron acceptors (Figure 3). All three electron acceptors oxidized MmcA as confirmed by the change in the spectral profile of the protein (Figure 3; Supplementary Figures 9 and 10). These data indicate the biochemical importance of MmcA for DSMR in *M. acetivorans*.

**Figure 3:**
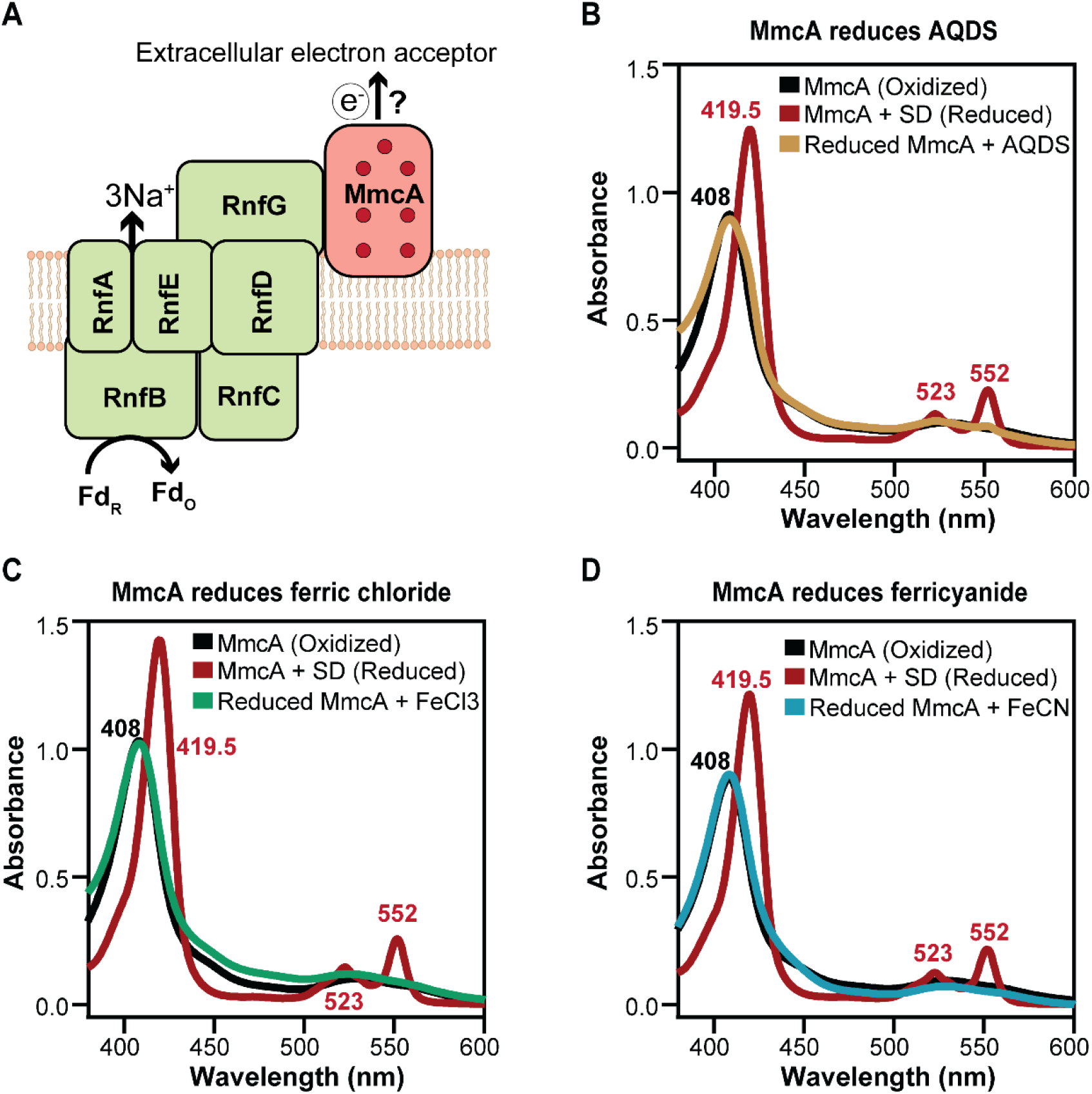
MmcA can donate electrons to extracellular electron acceptors. (**A**) Schematic of the MmcA-Rnf complex where the multiheme cytochrome MmcA mediates the final step in the transfer of electrons from the complex to extracellular electron acceptors. (**B-D**) Spectral analysis of MmcA mediated reduction of anthraquinone-2,6 disulfonate (AQDS) (**B**), ferric chloride (**C**), and ferricyanide (**D**) under anaerobic conditions. Addition of AQDS (**B**), ferric chloride (**C**), and ferricyanide (**D**) to MmcA reduced with sodium dithionite (SD; red) leads to the oxidation of MmcA (yellow in **B**; green in **C**; blue in **D**) as observed by the appearance of a characteristic Soret peak at 408 nm indicative of the oxidized protein (black).

### MmcA improves the rate of iron reduction by *M. acetivorans*

To test the role of MmcA during DSMR in *vivo*, we measured the rate of Fe(III) reduction with methanol as the electron donor in cell suspension assays (Fig. 4). Consistent with our expectations, the absence of *mmcA* slows down Fe(III) reduction rates by ∼30% *in vivo* (Fig.4B; Supplementary Figure 11). To test for polar effects of deleting *mmcA* on the chromosome, we measured the rate of iron reduction in the Δ*mmcA* mutant complemented with *mmcA* on a plasmid. Complementing *mmcA* in *trans* restored iron reduction rates to wildtype levels (Fig. 4B; Supplementary Figure 11). These data indicate that, even though MmcA is not the only mechanism for DSMR in *M. acetivorans,* it plays an important role during iron reduction.

**Figure 4:**
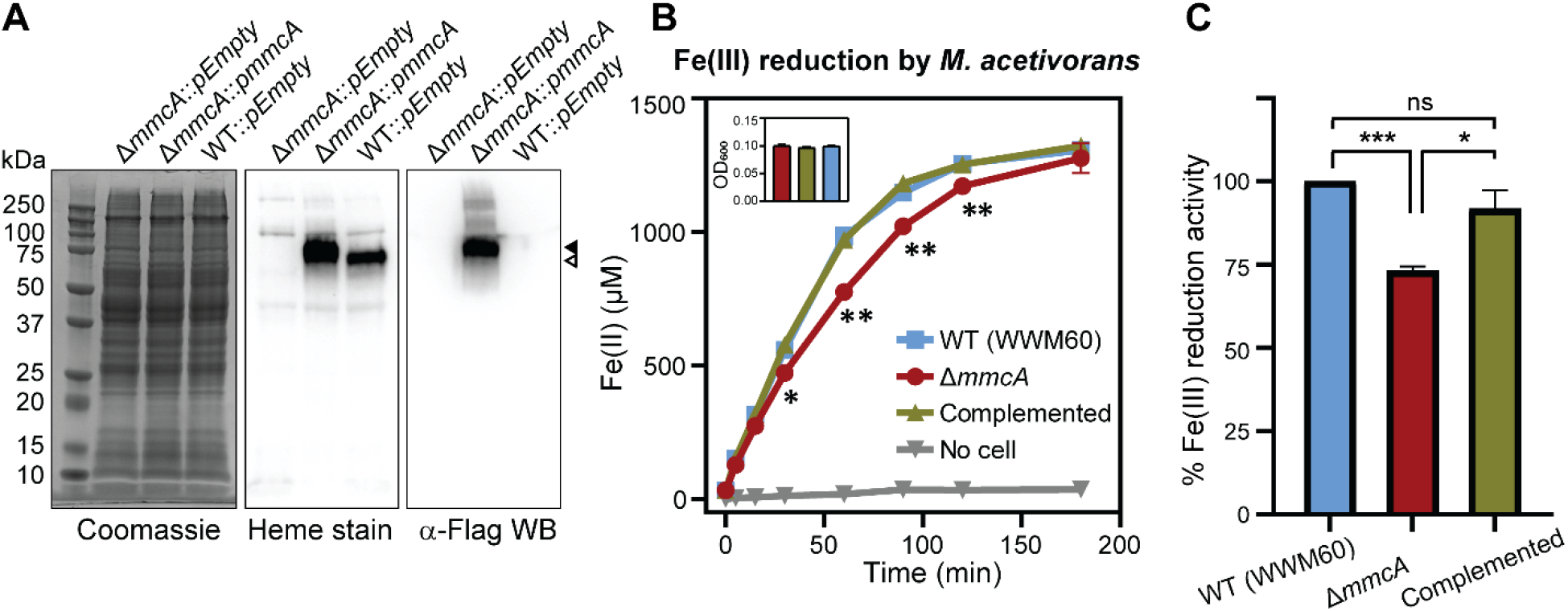
MmcA facilitates iron-reduction in *M. acetivorans*. (**A**) Coomassie, heme stain and Western blot (WB) with anti (α)-FLAG antibody of cell lysates obtained from a Δ*mmcA* mutant complemented with a plasmid encoding a C-terminal 3×FLAG tagged *mmcA* placed under the control of a tetracycline inducible promoter (Δ*mmcA*::*pmmcA*). The controls are either the Δ*mmcA* mutant or the parent strain (WWM60; wildtype or WT) with an empty vector (pDPG010; labelled here as *pEmpty*) as described before (21). Wildtype MmcA and C-terminal 3×FLAG tagged MmcA are depicted by open and filled arrows respectively. (**B**) The rate of Fe(III) reduction in 1 mL anaerobic cell suspensions with similar optical density (see inset) was determined by an increase in [Fe(II)] using the ferrozine assay. 100 µg/mL of tetracycline was used to induce expression. Error bars are means ± standard error of three technical replicates. (**C**) Rate of Fe(III) reduction by the Δ*mmcA* and Δ*mmcA*::*pmmcA* complementation strain were calculated as the percentage activity of the parent strain (WWM60) denoted as wild-type (WT). The data shown are average ± standard error from three independent experiments done with three technical replicates each. * p-value ≤ 0.05, ** p-value ≤ 0.01, *** p-value ≤ 0.001 and ns is p-value ≥ 0.05 respectively using a two-sided Student’s t-test.

### MmcA is reversibly redox-active between -100 and -450 mV versus SHE

We explored the redox behavior of MmcA using protein film voltammetry (PFV) on the meso-porous indium tin oxide electrode (ITO). MmcA formed a stable film on the ITO surface and was capable of exchanging electrons directly with the electrode, giving rise to an overlapping envelope of reversible signals spanning from -100 to -450 mV (Fig. 5A; Supplementary Figures 12 and 13). These signals, once deconvoluted and fit to Nernstian one-electron peaks, could be separated into seven reversible signals corresponding to the reduction and oxidation of the seven heme cofactors within MmcA, with uncertainties of approximately 10 mV. We determined the midpoint potentials of each heme by taking the average of the reduction and oxidation peak potentials for each redox pair (Fig. 5B). The low potential redox range of MmcA is similar to functionally analogous MHCs like MtrA and the MtrCAB complex (0 to -400 mV and 0 to -450 mV, respectively) in *Shewanella oneidensis* MR-1 (34, 35) and OmcZ or OmcS (−60 to −420 mV and −40 to −360 mV, respectively) in *Geobacter sulfurreducens* (36, 37). The redox-active range of MmcA suggests that it is capable of transferring electrons to Fe(III) (+0.3 to +0.4 V), AQDS (−0.185 V) and MP (−0.165 V) (Fig. 5B).

**Figure 5:**
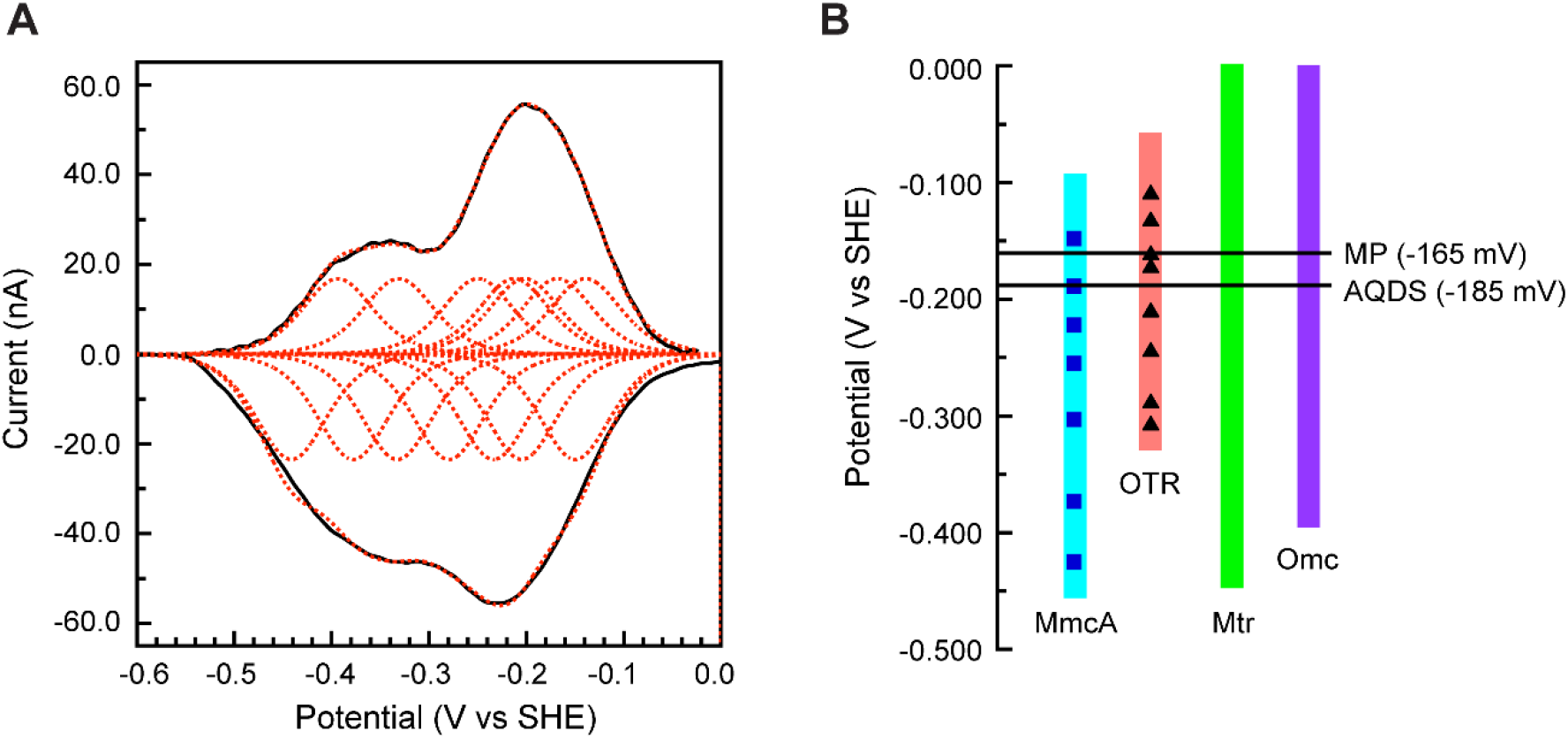
MmcA is redox active at a potential between -100 and -450 mV versus SHE. (**A**) Background-subtracted non-turnover MmcA voltammogram (black solid) with fitting of seven reversible redox couples (red dotted). Cyclic voltammetry (CV) was performed at pH 7.4, 10°C, and a scan rate of 20 mV/s. (**B**) Redox range of MmcA from *Methanosarcina acetivorans* and octaheme tetrathionate reductase (OTR) from *Shewanella oneidensis* (See Supplementary Figure 15). The average midpoint potentials (E_m_) of the seven heme centers in MmcA (blue squares) and eight heme centers in OTR (black triangles) are shown. The redox range of other multiheme *c*-type cytochromes like the metal reductase (Mtr) from *Shewanella oneidensis* (34, 35) and the outer membrane cytochromes (Omc) from *Geobacter sulfurreducens* (36, 37) as well as the midpoint potential of methanophenazine (MP) and anthraquinone-2,6-disulfonate (AQDS) are shown for reference.

### MmcA is related to the tetrathionate reductase (OTR) family of multiheme *c*-type cytochromes

MmcA is widespread in methanogens and anaerobic methane oxidizing archaea (ANME) within the Order *Methanosarcinales* but is otherwise absent in Archaea and Bacteria (26). To identify distant homologs of MmcA, we performed a BLAST search with the amino acid sequence of MmcA from *M. acetivorans* in Archaea and uncovered predicted octaheme tetrathionate reductases (OTR) in a few members of the *Methanosarcinales, Desulfurococcales* and *Archaeoglobales*. To understand how MmcA and OTR are related to each other, we generated a maximum-likelihood phylogenetic tree with the entire MmcA family, a few sequences that represent the diverse OTR family in Archaea and Bacteria and rooted this tree using distant MHCs like nitrate/nitrite reductases from Bacteria as an outgroup (Fig. 6A). In this analysis OTRs from Archaea cluster with OTRs from Bacteria rather than with MmcA, indicating that these genes were acquired independent of MmcA (Fig. 6A). Based on the topology of the tree, we can infer that the MmcA clade is distinct but closely related to OTRs. This observation is further corroborated by analyzing the genomic neighborhood of MmcA and OTR in methanogens (Fig. 6B). The genome neighborhood of the *mmcA* locus is universally conserved; it is always found in an operon with other genes of the Rnf complex. In contrast, in strains that encode *otr,* this gene is found in close proximity to a thermosome subunit and a biotin transporter (Fig. 6B).

**Figure 6:**
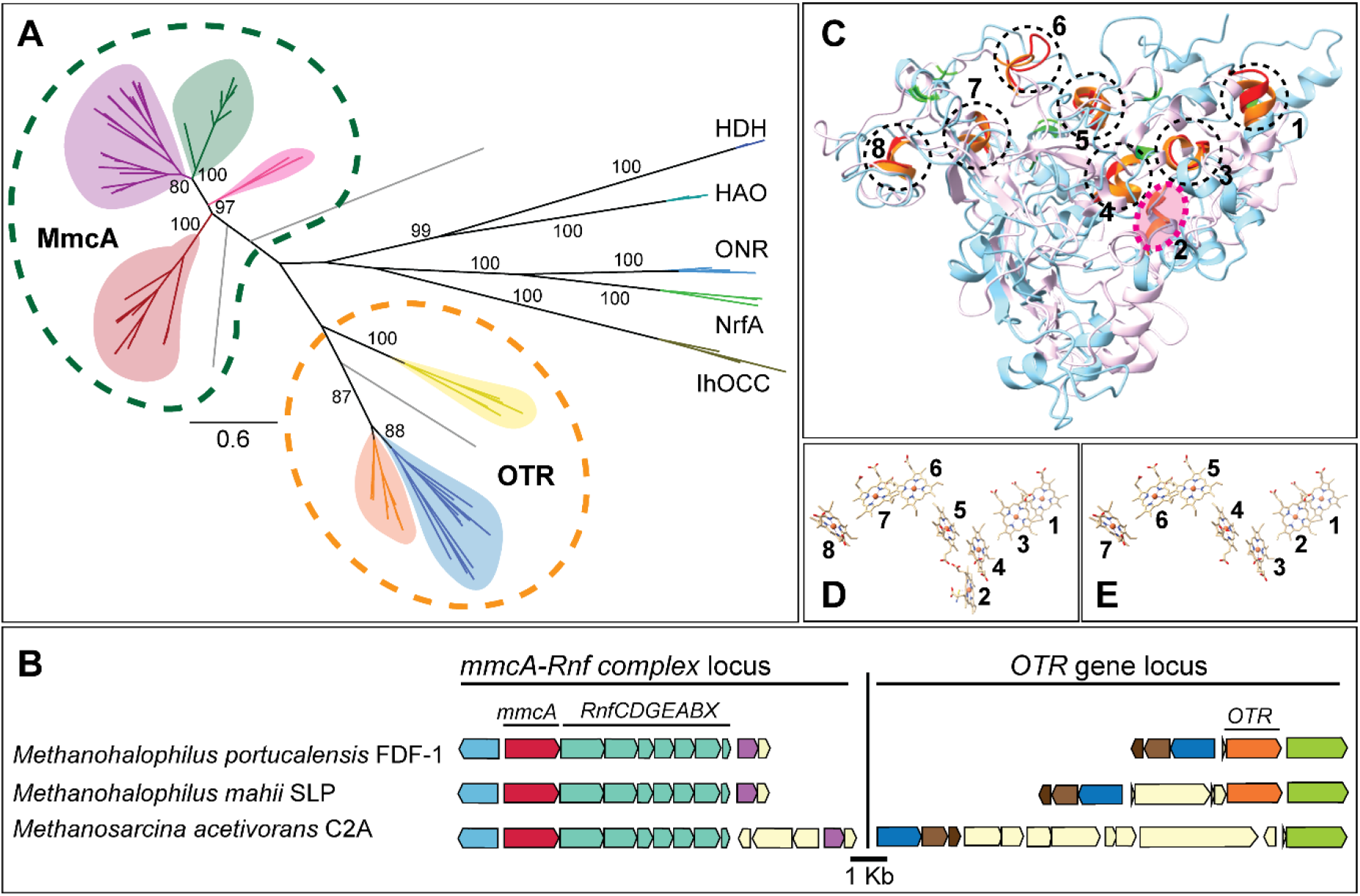
MmcA comprises a distinct clade of multiheme cytochromes related to the octaheme tetrathionate reductases (OTR). (**A**) Maximum-likelihood phylogenetic tree of MmcA and representatives of the OTR family. MmcA clades are colored by members of the Genus *Methanosarcina* (green), other members of the Family *Methanosarcinaceae* (purple), unspecified MAGs from the Order *Methanosarcinales* (pink), and anaerobic methane oxidizing archaea (ANME) (red). OTR clades are colored by members of the *Methanosarcinaceae* (orange), members of the *Desulfurococcales* and *Archaeoglobales* (yellow), and representatives from Bacteria (blue). A pentaheme nitrite reductase (NrfA), octaheme nitrite reductase (ONR and IhOCC), octaheme hydrazine dehydrogenase (HDH) and octaheme hydroxylamine oxidoreductase (HAO) derived from bacteria were used as the outgroup to root the tree. Bootstrap values of 80 and above are shown. (**B**) Chromosomal organization of genes surrounding the *mmcA* locus (red) and the *otr* locus (orange) in a few representative strains within the Family *Methanosarcinaceae.* Genes of the same color represent members of the same orthologous group. (**C**) Structural alignment of AlphaFold predicted model of signal-less MmcA (MA0658; 25-500 aa, cyan) to crystal structure of OTR from *Shewenella oneidensis* (PDB: 1SP3, pink). Heme binding motifs of OTR (orange) and MmcA (red) are shown. The histidine (His) ligand of the seven bis-His coordinated heme groups (1, 3–8) of OTR (light green) and seven heme groups (1–7) of MmcA (dark green) are shown. The second heme binding motif of OTR that is absent in MmcA is highlighted (pink circle). Arrangement of heme groups in the OTR crystal structure (**D**) and in the MmcA model (**E**).

Unlike MmcA, OTR is a respiratory enzyme of cryptic native function, best characterized as a reductase of tetrathionate, nitrite, and hydroxylamine in bacteria like *Shewanella oneidensis* (38, 39). All OTRs, including those derived from members of the *Methanosarcinales,* contain eight heme-binding motifs and structural studies show that the second heme group is ligated by a highly conserved Lys instead of His (38). Neither this heme-binding motif nor the Lys are conserved in MmcA (Supplementary Figure 14), which suggests that the function of MmcA has diverged from a canonical OTR with the loss of the Lys-bearing heme. Comparing an AlphaFold model of MmcA (MA0658) to OTR from *S. oneidensis* (1SP3.pdb) corroborates this view. The heme-binding motifs of MmcA are in structural alignment with seven of the eight heme binding motifs in OTR (Fig. 6C). As predicted by sequence alignment, the second heme binding motif in OTR, involved in catalysis, is missing in MmcA (Fig. 6D). All seven heme binding motifs in MmcA are within electron transfer range for the heme groups with a total head-to-head distance of approximately ∼55 Å between the first to seventh heme groups (Fig. 6E). Finally, the range of the redox potentials of the heme groups in MmcA (Fig. 5) is similar to that of OTR derived from *S. oneidensis* (Fig 5B, Supplementary Figure 15). Altogether, these data suggest that the MmcA clade of MHCs is distinct in form and function but closely related to the well-characterized OTR family of MHCs.

## Discussion

From a thermodynamic perspective, methanogens like *M. acetivorans* that possess an ETC and primarily use organic carbon compounds as electron donors for methanogenesis are well-suited for alternate energy conservation strategies. Switching to DSMR, say using methanol or acetate as an electron donor, in the presence of extracellular electron acceptors like ferric ions is much more thermodynamically favorable than disproportionation of these compounds to methane and carbon dioxide in these methanogens (see Supplementary Table 1). Here, we show that the presence of MmcA allows *M. acetivorans* to reconfigure its ETC in response to the environment as described above. During methanogenesis (i.e. in the absence of any extracellular electron acceptors) cells generate a heterodisulfide of CoM-S-S-CoB, which serves as the terminal electron acceptor for the ETC in *M. acetivorans* (3). Under these circumstances, MmcA transfer electrons from the Rnf complex to the membrane-bound electron carrier MP (Supplementary Figure 1; Figure 2), which can ultimately be used to reduce the terminal electron acceptor CoM-S-S-CoB. Notably, MmcA is only present in the Rnf complex of methane metabolizing archaea that use MP as a membrane-bound electron carrier, and is absent in Bacteria that use the Rnf complex to transfer electron between the ferredoxin and NAD pools (26). When *M. acetivorans* encounters extracellular electron acceptors, our data suggest that MmcA can directly interact with and reduce these electron acceptors (Figures 3 and 4). It has recently been proposed that the addition of Fe(III) to cultures of *M. acetivorans* enhances methanogenesis from acetate (13). Under these circumstances, MmcA would have to simultaneously channel electrons to MP and Fe(III). Based on the organization of the heme groups in the AlphaFold model of MmcA it is feasible to chart a path for electron flow through MmcA so that cells can perform DSMR and methanogenesis at once (Supplementary Figure 16). This proposed path of electron flow through MmcA is potentially beneficial as a small amount of methane production seems vital for the growth of methanogens even under non-methanogenic conditions likely due to an auxiliary role for CoM-S-S-CoB that is yet to be identified (40).

Electroactive microorganisms can use the same set of MHCs to transport electrons in or out of the cell (41, 42). Accordingly, recent studies have suggested that MmcA might facilitate electron uptake from metallic iron or a partner bacterium via direct interspecies electron transfer to fuel methanogenesis (19, 20). These data are consistent with our electrochemical observations, which show that MmcA can reversibly exchange electrons with the electrode during PFV. Additionally, even though MmcA is important for EET in *M. acetivorans,* we cannot rule out the possibility that *M. acetivorans* encodes additional routes for EET for a number of reasons. First, the Δ*mmcA* mutant of *M. acetivorans* can still reduce Fe(III), albeit at lower rates (Figure 4). Second, *M. barkeri* does not encode *mmcA* or any other cyt *c* in its genome yet can perform DSMR (6, 9–11). While a hydrogenase-dependent EET pathway is plausible for *M. barkeri,* it is unlikely for *M. acetivorans* as this strain does not exhibit any hydrogenase activity under a wide range of laboratory conditions tested so far (43, 44). It is also worth noting that a hydrogenase-independent pathway for extracellular electron uptake has been demonstrated for a mutant of *Methanococcus maripaludis* lacking all catabolic and anabolic hydrogenases (45). An unbiased transposon mutagenesis screen in the Δ*mmcA* mutant of *M. acetivorans* coupled with high-throughput assays of the mutant library for Fe(III) reduction might help identify additional pathways for EET in the future.

The spatial organization of MmcA-mediated EET in *M. acetivorans* differs from other microorganisms. In Gram-negative bacteria like *S. oneidensis* MR-1, EET is coordinated by three distinct electron transfer steps: i) from the inner membrane to the periplasmic space, ii) from the periplasm to the outer membrane and iii) from the cell surface to the extracellular electron acceptor(s) (46, 47). Each of these three steps is mediated by a distinct set of MHCs that are appropriately located and have unique electrochemical properties (46, 47). A similar process occurs in Gram-positive bacteria like *Thermincola potens* where the final step of EET is mediated by MHCs anchored in the cell wall (48). In contrast, most methanogenic archaea like *M. acetivorans* lacks a cell wall altogether and their cell envelope comprises of an inner membrane and a crystalline proteinaceous surface layer (S-layer) (49). Cell aggregates of *M. acetivorans* are embedded in an extracellular matrix composed of methanochondroitin that is practically absent in planktonic cells used in the study (50). The hexagonal crystal packing of the S-layer forms three different pores of which one, the primary pore, is large enough to allow the passage of chelated metal ions and humic acid analogs like AQDS into the pseudo-periplasm (51). As a result, it is likely that MmcA could interact with extracellular electron acceptors in the pseudo-periplasmic space. Whether MmcA could directly interact with metal minerals or other cells to support direct electron transfers is currently unclear. However, it is plausible that redox-active species like soluble Fe(III) and AQDS could shuttle electrons between MmcA and minerals or cells. Microbes are also known to produce small redox-active molecules such as flavin and phenazine to shuttle electrons during EET (52, 53). However, it is currently unknown if *M. acetivorans* produces any such molecules that can act as a shuttle for EET.

In conclusion, we show that MmcA controls the flow and path of electrons through the respiratory chain that, in turn, tunes the carbon flux through methane in methanogens like *M. acetivorans*. Based on its cellular role, MmcA has the potential to be a focal point of engineering efforts to divert carbon flux away from methane in methanogens and curb global emissions of this potent greenhouse gas.

## Materials and Methods

### Growth Medium

*M. acetivorans* strains (Supplementary Table 2) were grown in single-cell morphology (50) at 37°C without shaking in bicarbonate-buffered high-salt (HS) liquid medium containing either methanol or trimethylamine (TMA) as the carbon and energy substrate with N_2_/CO_2_ (80/20) in the headspace. Puromycin (RPI, Mount Prospect, IL) was added to a final concentration of 2 µg/mL from a sterile, anaerobic stock solution to select for *M. acetivorans* strains with the *mmcA-*expression plasmid encoding a puromycin-resistance gene (*pac)*. Anaerobic, sterile stocks of tetracycline hydrochloride in deionized water were prepared fresh before use and added to a final concentration of 100 µg/mL to induce the expression of MmcA from the tetracycline-inducible promoter as described previously in (54). Cell cultures with a volume of up to 10 mL were grown in Balch tubes and larger volume cultures were grown in anaerobic bottles.

### MmcA purification

MmcA was affinity purified from 4 L of late-exponential phase culture of DDN039 grown in HS media with 100 mM TMA at 37°C. Cells were harvested by centrifugation (6000 x g) for 20 minutes at 4°C, the supernatant was discarded, and the cell pellets were stored at -80°C. All steps of protein purification were performed under aerobic conditions. 2 U/mL DNase-I (to reduce the viscosity of the suspension) and 1 mM Phenylmethylsulfonyl fluoride (PMSF) (to inhibit protease activity) was added to 20 mL of hypotonic lysis buffer (50 mM Tris-HCl, pH = 7.4) used to resuspend the cell pellet. The cell suspension was kept on ice for 45 min with intermittent mixing using a pipette to lyse the cells. Upon complete lysis, sodium chloride was added from a 5 M stock solution to a final concentration of 150 mM to the cell lysate. The lysate was clarified by centrifugation at 10,000 x g for 20 minutes at 4°C and the supernatant was separated into the soluble and membrane fractions by high-speed ultracentrifugation at 100,000 x g for 1 hour at 4°C. The membrane pellets were solubilized in 4 mL TBS buffer (50 mM Tris-HCl, 150 mM NaCl, pH = 7.4) with 2% Triton X-100 (Sigma-Aldrich, St Louis, MO, USA). The solubilized membrane-fraction was loaded on a column containing 1 mL anti-DYKDDDDK (Flag) G1 affinity resin (50% suspension; GenScript, Piscataway, NJ, USA) pre-equilibrated with 3 bed volumes of TBS buffer. Five washes with 2 mL of TBS buffer were performed before the protein was eluted using competitive elution buffer (300 ug/mL Flag peptide in TBS buffer). To elute the protein, three times the bed volume (i.e., 1.5 mL) of elution buffer was added to the column and one volume (500 µL) of elute was collected right away. The column was capped and incubated at room temperature for 30 min before collecting the rest of the eluate. The elutes were quantified using Bradford reagent (Sigma-Aldrich, St Louis, MO, USA) with BSA (bovine serum albumin) as the standard following the manufacturer’s instructions and saved at -80°C.

### Proteolytic digestion of MmcA and LC-MS analysis

MmcA was digested with trypsin or chymotrypsin in solution for LC-MS analysis as previously described (55). Briefly, 20 µL of purified MmcA at a concentration of 2 mg/mL (total 40 µg) was mixed with 9.6 mg of urea (8M) in a sterile microfuge tube and incubated for 1 hour at room temperature for protein denaturation. The MmcA-Urea mix was diluted 10-fold by adding 180 µL freshly prepared 50 mM ammonium bicarbonate solution. Two aliquots of 100 µL were transferred into new tubes and digested either with a sequencing-grade trypsin (Promega, Madison, WI, USA) or chymotrypsin (Promega, Madison, WI, USA) per manufacturer’s instructions. For trypsin digestion, 2.5 µL of 0.4 µg/µL trypsin (1 ug) was added to 100 µL protein solution (1:20, enzyme to protein ratio) and incubated at 37°C overnight (*ca.* 16 hours). Similarly, 2 µL of 0.5 µg/µL chymotrypsin (1 µg) was added to another 100 µL protein solution (1:20, enzyme to protein ratio) and incubated at 25°C overnight (18 hours). 75 µL of the overnight-digest were transferred to a clean microfuge tube and submitted for MS analysis (QB3/Chemistry Mass Spectrometry Facility, UC Berkeley). Protein digests were also confirmed by running the remaining 25 µL sample on SDS-PAGE gels followed by Coomassie staining.

### UV-visible (vis) absorption spectroscopy with MmcA

All UV-vis spectroscopy was performed at room temperature with a Shimadzu 1900i (Shimadzu, Torrance, CA, USA) kept inside an anaerobic chamber (97% N_2_, and 3% H_2_; Coy Laboratory, Grass Lake, USA). Unless specified, all assays were conducted with 2.5-3.5 µM MmcA in 50 µL of assay buffer (50 mM Tris-HCl, 150 mM NaCl, 2% glycerol, pH = 7.4). Stock solutions of 1 mM and 10 mM sodium dithionite were prepared in deionized water. 2-hydroxyphenazine was custom synthesized (AstaTech Inc., Bristol, PA) and a 20 mM stock solution was prepared in 100% ethanol. A 10 mM stock of Anthraquinone-2,6-disulfonate (AQDS) was prepared in deionized water as described before (42). The solution was heated at 60°C until AQDS was completely dissolved (∼10 min), cooled down to room temperature and the pH was adjusted to 7.0. Stocks of 10 mM ferric chloride (FeCl_3_) and potassium ferricyanide (K_3_[Fe(CN)_6_]) were prepared with deionized water. All assay components (buffer, protein, chemicals) were kept in the anaerobic chamber in small volumes (30-50 µL) at least 2 hours prior to the assay and were confirmed to be anaerobic by using the redox dye resazurin (0.0001 % w/v) and testing for a color change from colorless to pink after 10 mins.

### Pyridine hemochrome assay

This assay was performed as described before (56, 57). Briefly, a 0.2 M NaOH with 40% pyridine solution was made fresh using a 1 M NaOH stock and 100% pyridine solution (Sigma Aldrich, St. Louis, MO). 5 µL (i.e., 1/200) of 0.1 M potassium ferricyanide stock solution was added 495 µL of the aforementioned NaOH + pyridine mix to generate the pyridine hemochrome assay solution. 50 µL of the assay solution was mixed with 50 µL of TBS buffer (50 mM Tris-HCl, 150 mM NaCl, pH = 7.4) and used as a blank. Next, 50 µL of the assay solution was mixed with 50 µL of MmcA in TBS buffer, and UV-vis scans were immediately performed using a Shimadzu 1900i (Shimadzu, Torrance, CA, USA) to record the oxidized spectra. A 10 mM stock solution of sodium dithionate was added to the protein assay mixture and UV-vis scans were performed using a Shimadzu 1900i (Shimadzu, Torrance, CA, USA) to record the fully reduced pyridine hemochrome spectra.

### Cell suspension assays

Cell suspension assays were performed in an anaerobic chamber (97% N_2_, and 3% H_2_; Coy Laboratory, Grass Lake, USA) at room temperature as previously described for bacterial cells (58) with some modifications. Assays were performed with an *M. acetivorans* mutant lacking the chromosomal copy of the *mmcA* locus and expressing *mmcA* from an inducible promoter on a plasmid and a control strain containing the plasmid pDPG010 described previously (21). All strains were grown in HS medium with 125 mM methanol, 2 µg/mL puromycin and *mmcA* expression was induced by adding tetracycline to a final concentration of 100 µg/mL. Cells were harvested in mid-exponential phase (OD_600_ = 0.4 – 0.6) by centrifugation in the anaerobic chamber. The cell pellet was resuspended in anaerobic high-salt PIPES buffer (50 mM PIPES, 400 mM NaCl, 13 mM KCl, 54 mM MgCl_2_, and 2 mM CaCl_2_, pH 6.8) containing 5 mM methanol and washed three times. At the end of the third wash, cells were resuspended in anaerobic high-salt PIPES buffer containing 5 mM methanol and supplemented with a freshly prepared 1:1 Fe(III)-nitrilotriacetic acid (NTA) mix using an anaerobic 0.4 M Fe(III) chloride and an anaerobic 0.8 M NTA stock solutions to final concentration of 1 mM Fe(III) chloride and 2 mM of NTA. Fe(III) reduction was monitored by sampling the suspension at different time points and measuring the Fe(II) concentration by using the ferrozine assay as described previously (59).

### Electrochemistry

Protein film voltammetry (PFV) experiments were carried out using a three-electrode cell configuration with the cell thermostated at 10 °C and housed inside a nitrogen-filled MBraun Labmaster glovebox (residual O_2_ < 1 ppm). The reference electrode was a saturated calomel electrode (SCE), and the counter electrode a platinum wire. The working electrode was a meso-porous indium tin oxide (ITO) electrode, prepared according to reported procedures (60). Briefly, a pyrolytic graphite edge (PGE) electrode (3 mm diameter) was polished and sonicated in water. ITO nanoparticles (Sigma, < 50 nm) were then deposited on the PGE surface by electrophoretic deposition. The PGE electrode was submerged in an acetone solution (20 mL) of I_2_ (0.01 g) and ITO (0.02 g), and a potential of 10 V was applied for 6 min using a graphite rod as the auxiliary electrode that was held approximately 1 cm away. The ITO electrode was then thoroughly rinsed with water and dried prior to use. To deposit the protein, a 3 μL aliquot of the protein solution (100 μM MmcA or 55 μM OTR, the latter prepared as previously described (61) was placed on the electrode for 3 min. Excess protein solution was then removed by rinsing with cold buffer, and the electrode was immediately placed into the electrolyte buffer, which contained 10 mM MES, 10 mM MOPS, 10 mM TAPS, 10 mM CHES, 10 mM HEPES, 10 mM CAPS, and 200 mM NaCl (pH 7.4). Cyclic voltammograms (CVs) were collected using the GPES software package (Ecochemie) that was connected to a PGSTAT30 AutoLab potentiostat (Ecochemie). All PFV data were analyzed using the qSOAS package (62), through which background electrode capacitance was subtracted, and data were filtered to remove electrical noise. Deconvolution of the redox feature was achieved within qSOAS using procedures reported previously, where the redox stoichiometry of all 7 heme cofactors was set to 1.0 (n = 1.0).

### Bioinformatics analyses

MmcA homologs were extracted from the NCBI non-redundant protein database using the MmcA (MA0658) protein sequence from *M. acetivorans* as the query and Archaea as the search database. Alignments and tree building were conducted in Geneious Prime 2023.0.3 (https://www.geneious.com). Any partial sequences (<375 aa) were discarded. The sequences were aligned using MUSCLE with default parameters. Maximum-likelihood tree of MmcA was generated using RAxML (Protein Model - GAMMA BLOSUM62; Algorithm - Rapid Bootstrapping and search for best-scoring ML tree; Number of starting trees or bootstrap replicates - 100; Parsimony random seed - 1). Gene ortholog neighborhood analysis was performed on Integrated Microbial Genomes and Microbiomes using bidirectional best BLAST hits for the gene of interest (63). For structural alignment and model building, an AlphaFold2 prediction of MmcA (64) and its closest available crystal structure i.e., 1SP3 for OTR from *Shewanella oneidensis* (SO4144) (38), were docked using matchmaker tools and 1SP3 as a reference structure in ChimeraX (65).

## Funding Information

The authors acknowledge funding from the ‘New Tools for Advancing Model Systems in Aquatic Symbiosis’ program from the Gordon and Betty Moore Foundation (GBMF#9324 to D.D.N. and D.G). The authors also acknowledge the support of the National Institutes of Health/National Institute of General Medicine (R35-GM136294 to S.J.E.). D.D.N. would also like to acknowledge funding from the Searle Scholars Program sponsored by the Kinship Foundation, the Rose Hills Innovator Grant, the Beckman Young Investigator Award sponsored by the Arnold and Mabel Beckman Foundation and the Packard Fellowship in Science and Engineering sponsored by the David and Lucille Packard Foundation. D.D.N is a Chan-Zuckerberg Biohub – San Francisco Investigator. The funders had no role in the conceptualization and writing of this manuscript or the decision to submit the work for publication.

## Author contributions

D.G. contributed to conceptualization, data curation, formal analysis, methodology, and writing. K.C. contributed to data curation, formal analysis, methodology, and writing. S.J.E. contributed to conceptualization, data curation, formal analysis, supervision, funding acquisition, methodology, and writing. D.D.N contributed to conceptualization, data curation, formal analysis, supervision, funding acquisition, project administration, methodology, and writing.

## Acknowledgements

We would like to acknowledge Dr. Anthony Iavarone for LC/MS analyses of peptide fragments of MmcA, Prof. Donald Rio for access to an ultracentrifuge for protein purification, Dr. Catarina Paquete for the donation of the *Shewanella oneidensis* strain used to produce OTR, Daniel Tekverk for producing OTR, and to members of the Nayak lab for their feedback and input.

## Competing Interests

The authors do not declare any competing interests.

## Supplementary Material

**Supplementary Figure 1:**
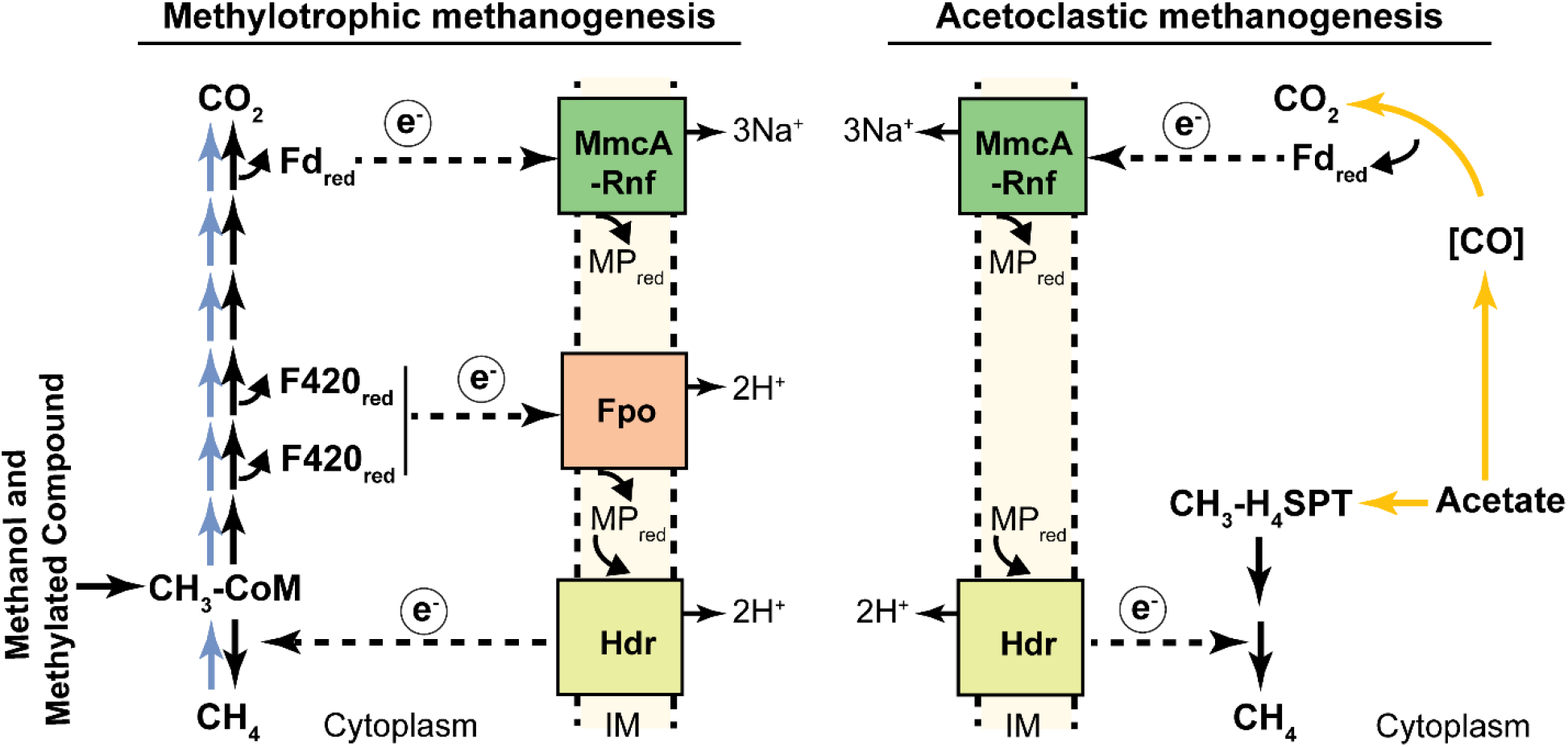
Role of MmcA-Rnf complex in methane metabolism. During growth on methylated compounds or methylotrophic methanogenesis (black arrows on left) and on acetate or acetoclastic methanogenesis (yellow arrows on right) in cytochrome-containing methanogens like *Methanosarcina acetivorans*, reduced ferredoxin is generated. It is hypothesized that anaerobic methanotrophic archaea (ANME) also produce reduced-ferredoxin as a result of anaerobic oxidation of methane (blue arrows on left). In all of these scenarios, electrons from reduced ferredoxin enter the electron transport chain through the MmcA-containing **<u>R</u>**hodobacter **<u>n</u>**itrogen **f**ixation complex (MmcA-Rnf). Reduced Ferredoxin (Fd_red_), reduced coenzyme F420 (F420_red_), inner cytoplasmic membrane (IM), sodium gradient (Na^+^), proton gradient (H^+^), F420 dehydrogenase complex (Fpo), heterodisulfide reductase complex (Hdr), tetrahydrosarcinapterin (H4SPT) and coenzyme M (CoM).

**Supplementary Figure 2:**
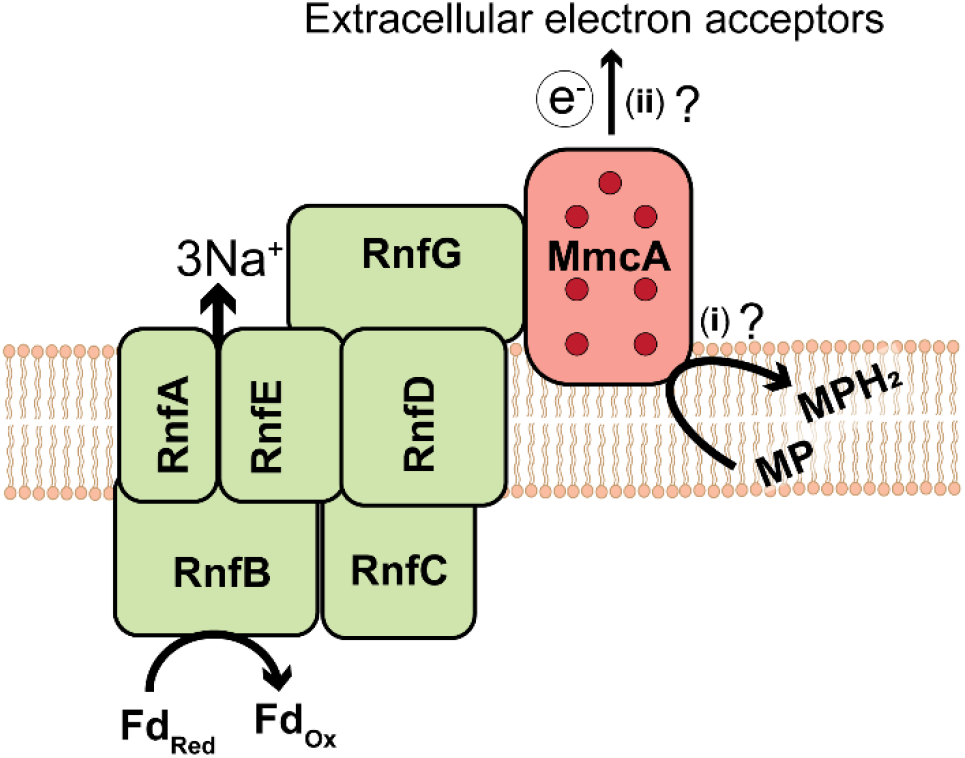
Role of MmcA in the Rnf complex of methane metabolizing archaea. The two proposed roles of MmcA are: (i) to transfer electrons from Rnf to methanophenazine (MP) (Li et al., 2006; Wang et al., 2011) and (ii) to transfer electrons to extracellular electron acceptors such as anthraquinone-2,6-disulfonate (AQDS) (Holmes et al., 2019), are shown. Red circles on MmcA represent heme binding motifs.

**Supplementary Figure 3:**
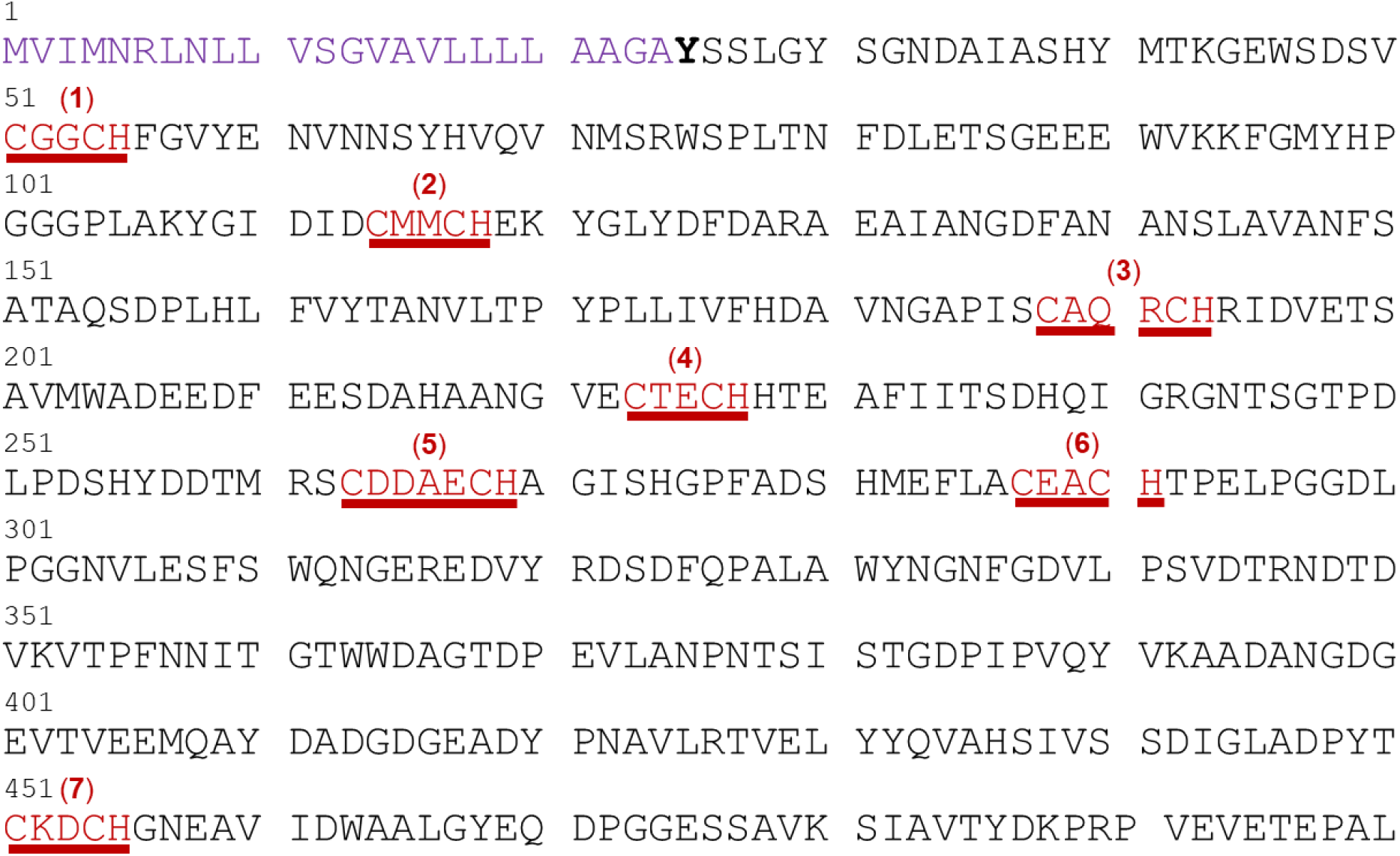
Amino acid (aa) sequence of the MmcA open reading frame (ORF) from *Methanosarcina acetivorans*. The 500 aa long apo-MmcA (encoded by MA0658) contains, a N-terminal Sec signal peptide (1-24 aa; shown in purple), heme binding motifs (underlined in red). MmcA contains five canonical heme binding motifs (CXXCH; 1, 2, 4, 6 and 7) and two heme binding motifs with extended gap between cysteine residues (3 and 5). Holo-MmcA is predicted to be 476 aa long protein (the predicted N-terminal amino acid in Bold) with seven covalently attached hemes.

**Supplementary Figure 4:**
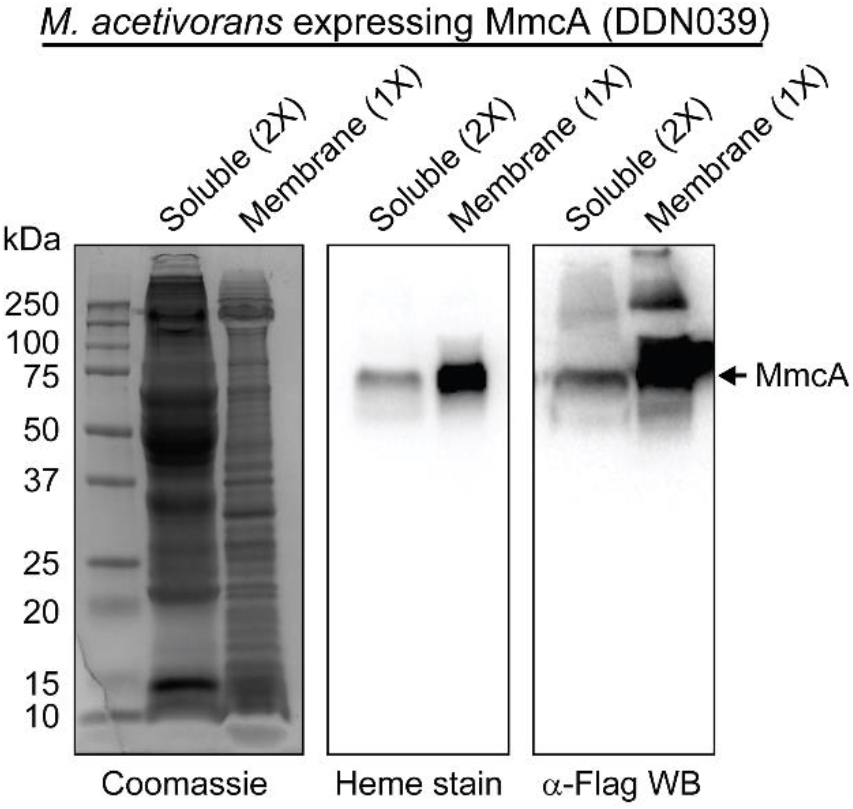
MmcA is primarily a membrane-associated protein. Coomassie, heme staining and Western blot (WB) with anti (α)-Flag antibody of the soluble and membrane fractions of *M. acetivorans* with the C-terminal 3×FLAG tagged MmcA-overexpression vector (DDN039). Twice the amount of the soluble fraction (2X, ∼40 µg) compared to the membrane fraction (1X, ∼20 µg) was loaded to visualize the small amount of MmcA in the soluble fraction. Based on the intensity of the heme stain and the Western blot, we can conclude that MmcA is primarily present in the membrane-fraction of cells.

**Supplementary Figure 5:**
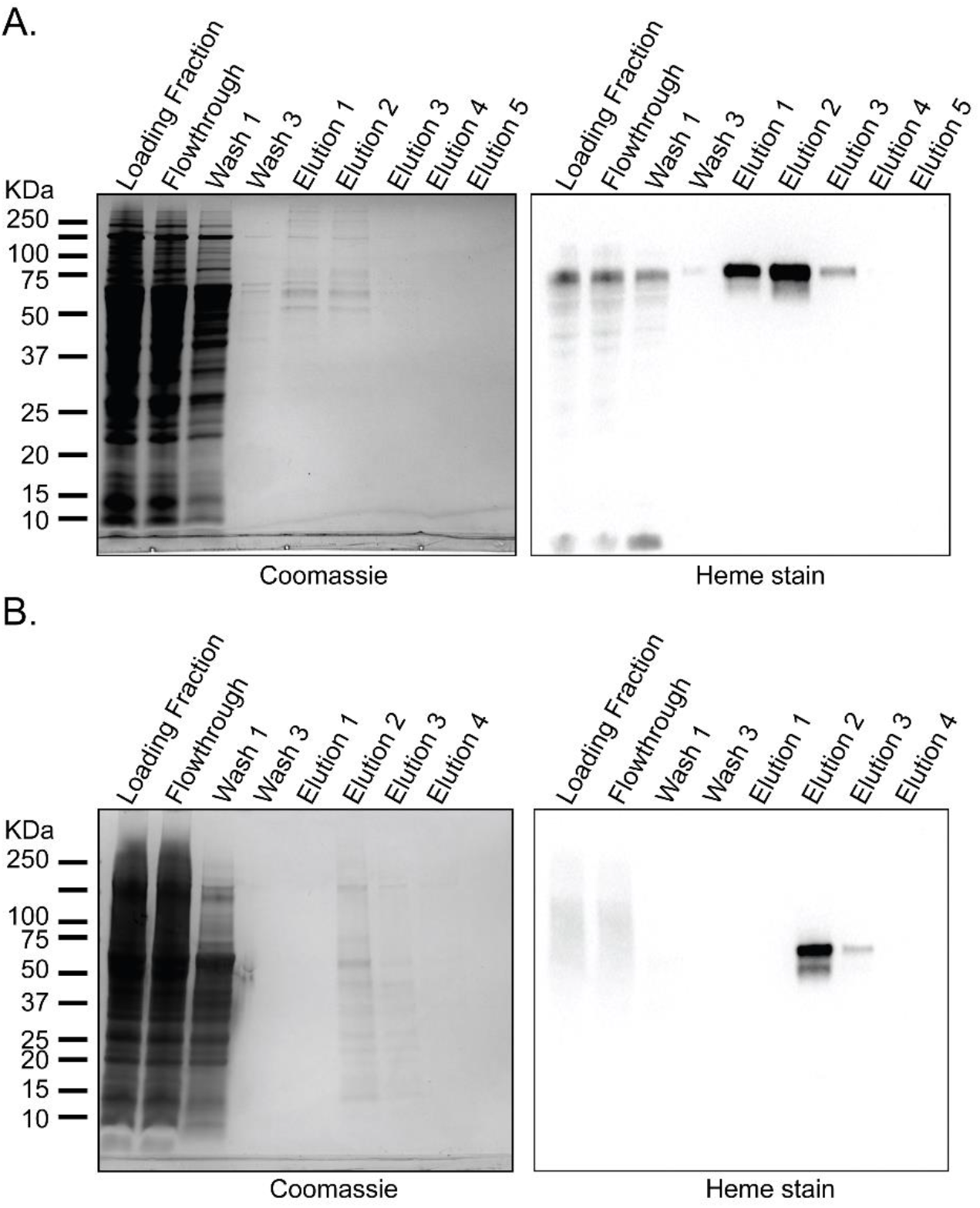
Affinity purification of C-terminal twin-Strep and 3×FLAG tagged MmcA using streptactin resin. Coomassie and heme staining of different fractions during MmcA purification using streptactin resin with **(A)** phosphate buffer (pH 8) and **(B)** Tris-buffer (pH 7.4). Trace amounts of MmcA was detected by heme staining.

**Supplementary Figure 6:**
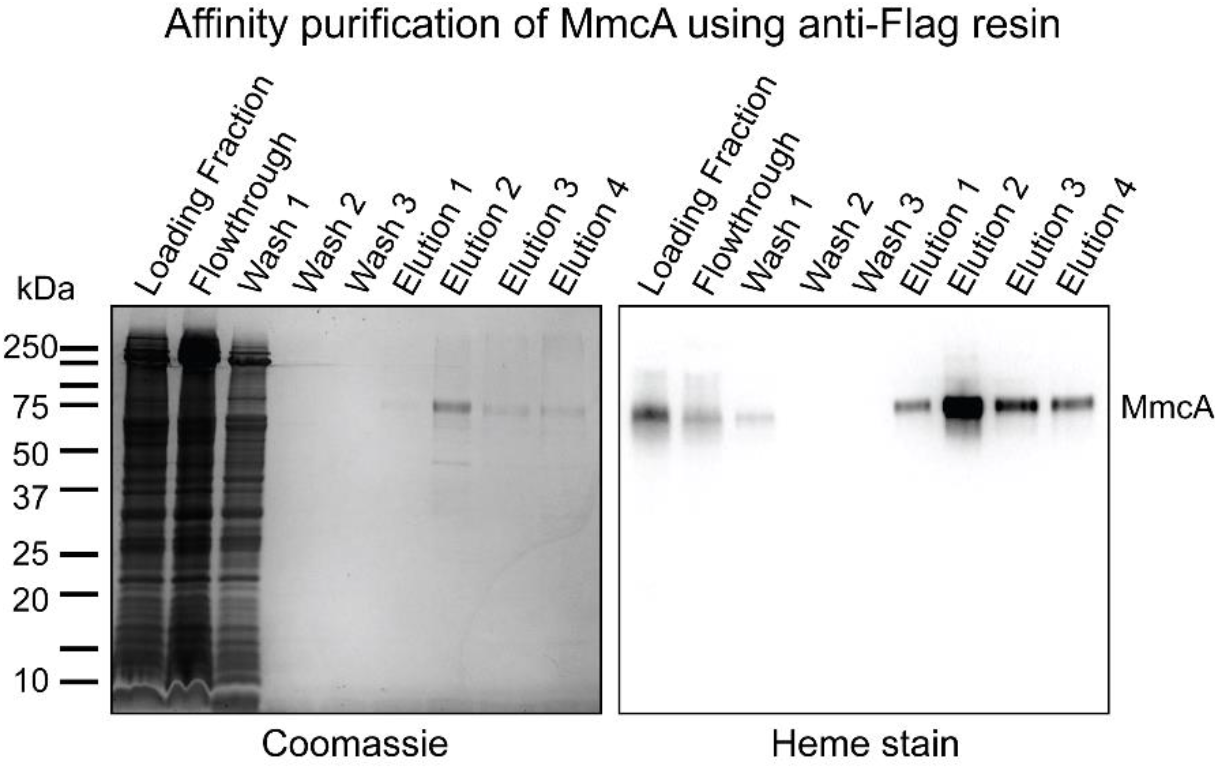
Affinity purification of C-terminal twin-Strep and 3×FLAG tagged MmcA using anti-Flag resin. Coomassie and heme staining of different fractions during MmcA purification using anti-Flag resin and Tris-buffer (50 mM Tris-HCl, 150 mM sodium chloride, pH = 7.4). A distinct MmcA band was visible in both Coomassie and heme staining.

**Supplementary Figure 7:**
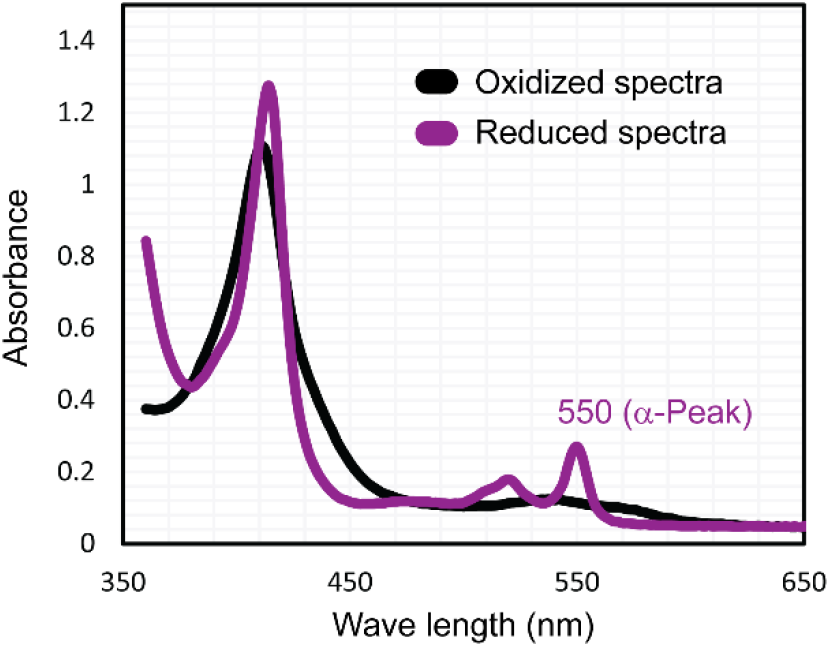
Pyridine hemochrome assay shows 550 nm α-Peak for MmcA. Oxidized (black) and reduced (purple) spectra of MmcA in pyridine hemochrome assay.

**Supplementary Figure 8:**
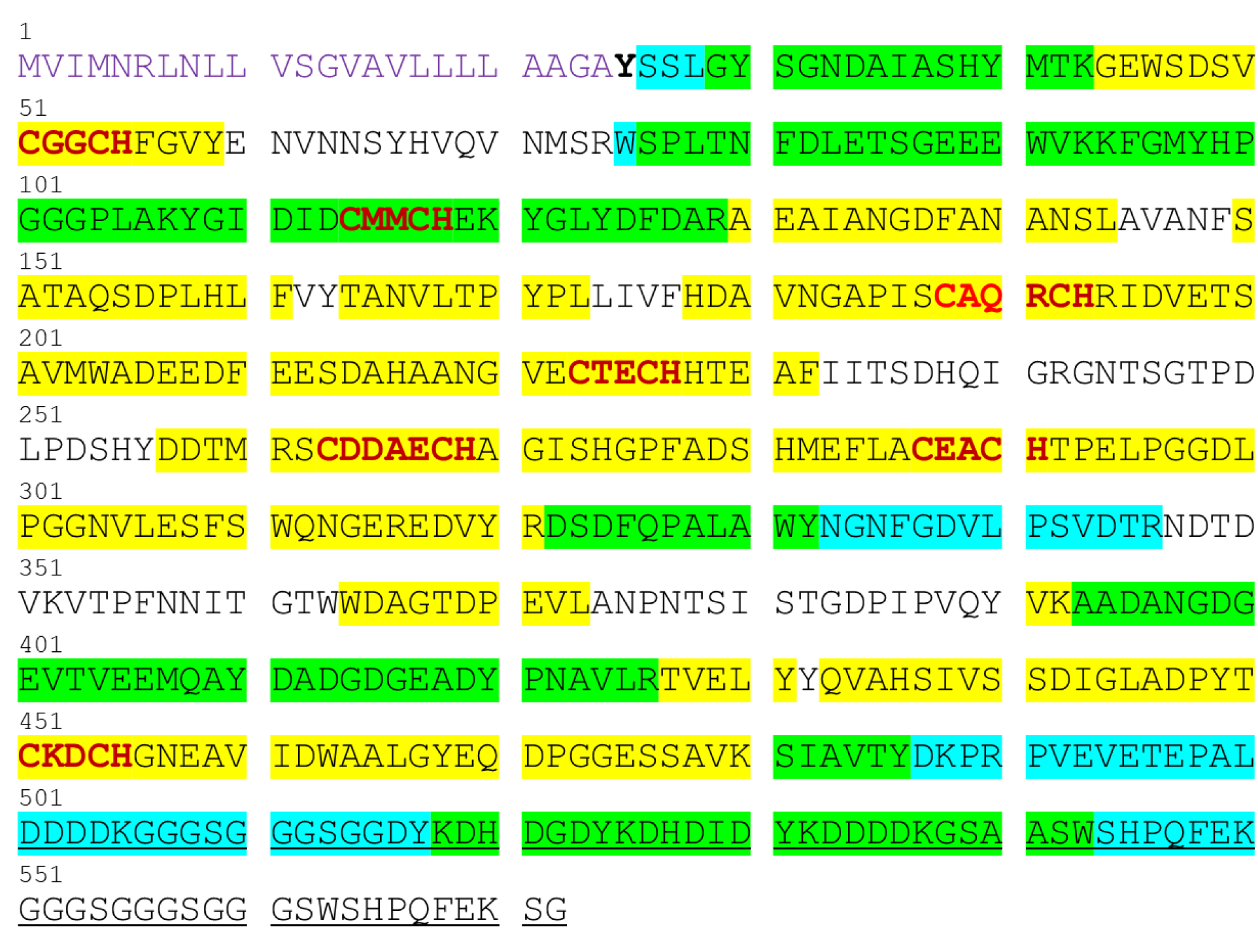
Mass spectrometry analyses of purified MmcA. Amino acid (aa) sequence of MmcA (encoded by MA0658) with tandem affinity purification (TAP) tag (underlined), a predicted N-terminal Sec signal peptide (1-24 aa; shown in purple), heme binding motifs (bold red) and the predicted N-terminal amino acid after Sec signal processing (bold black) are shown. Peptides detected by LC/MS analyses of purified MmcA digested with either trypsin or chymotrypsin are highlighted in cyan and yellow respectively. Peptides detected under both conditions are highlighted in green.

**Supplementary Figure 9:**
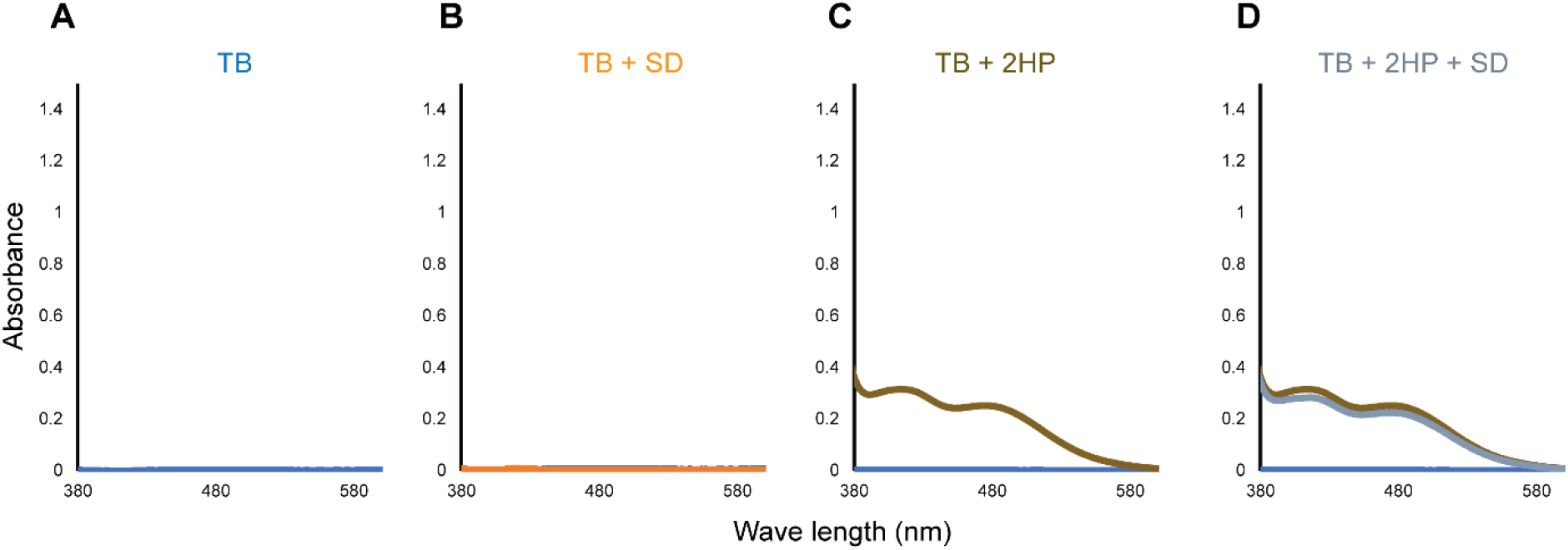
Protein-free controls for MmcA assays with 2-hydroxyphenazine. (**A)** Spectra observed for Tris buffer (TB), **(B)** TB plus sodium dithionite (SD), (**C**) TB plus 2-hydroxyphenazine (2HP), and **(D)** TB plus 2HP and SD.

**Supplementary Figure 10:**
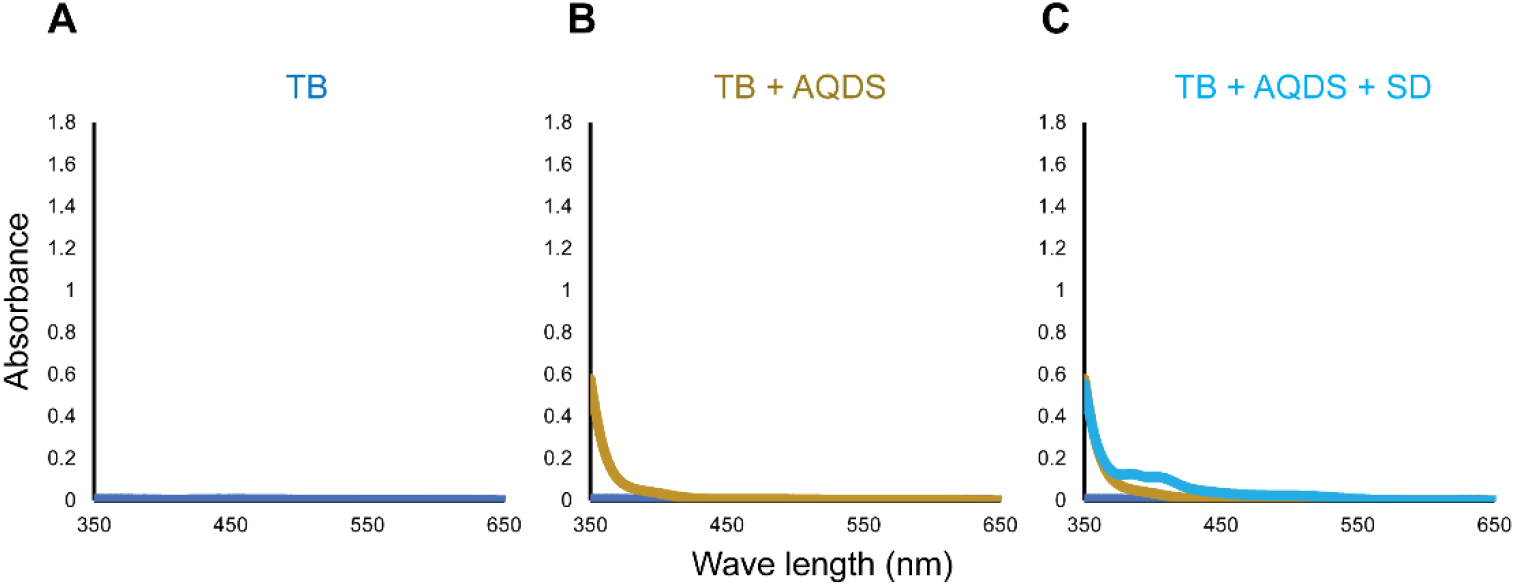
Protein-free controls for MmcA assays with anthraquinone-2, 6 disulfonate (AQDS). **(A)** Spectra observed for Tris buffer (TB), **(B)** TB plus AQDS, **(C)** TB plus AQDS and sodium dithionite (SD).

**Supplementary Figure 11:**
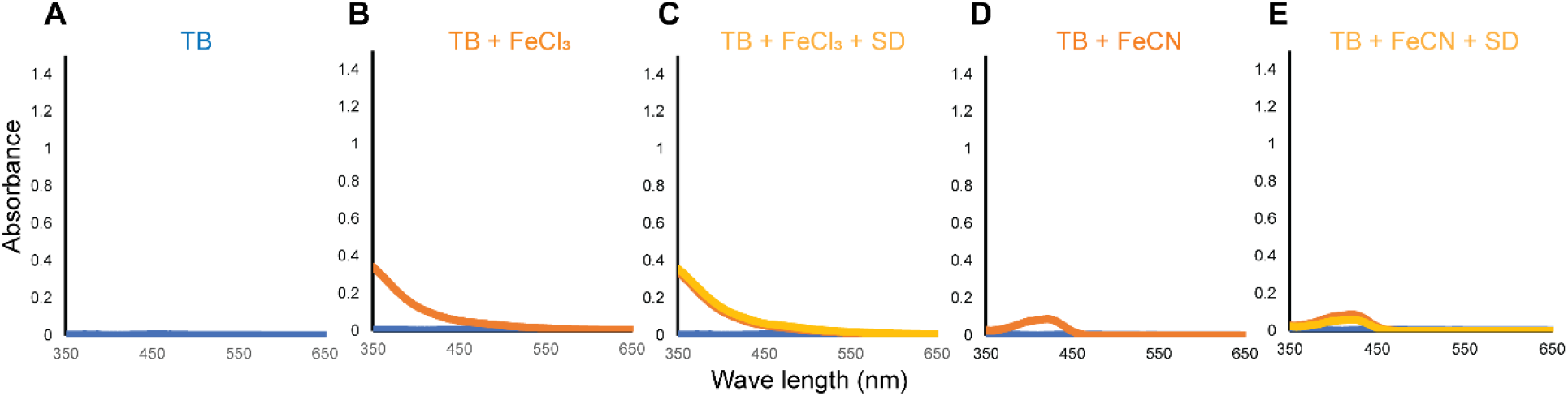
Protein-free controls for MmcA assays with ferric chloride and ferricyanide in UV-visible spectral analyses. **(A)** Spectra observed for Tris buffer (TB), **(B)** TB plus ferric chloride (FeCl_3_), **(C)** TB plus FeCl_3_ and sodium dithionite (SD), **(D)** TB plus ferricyanide (FeCN), and **(E)** TB plus FeCN and SD.

**Supplementary Figure 12:**
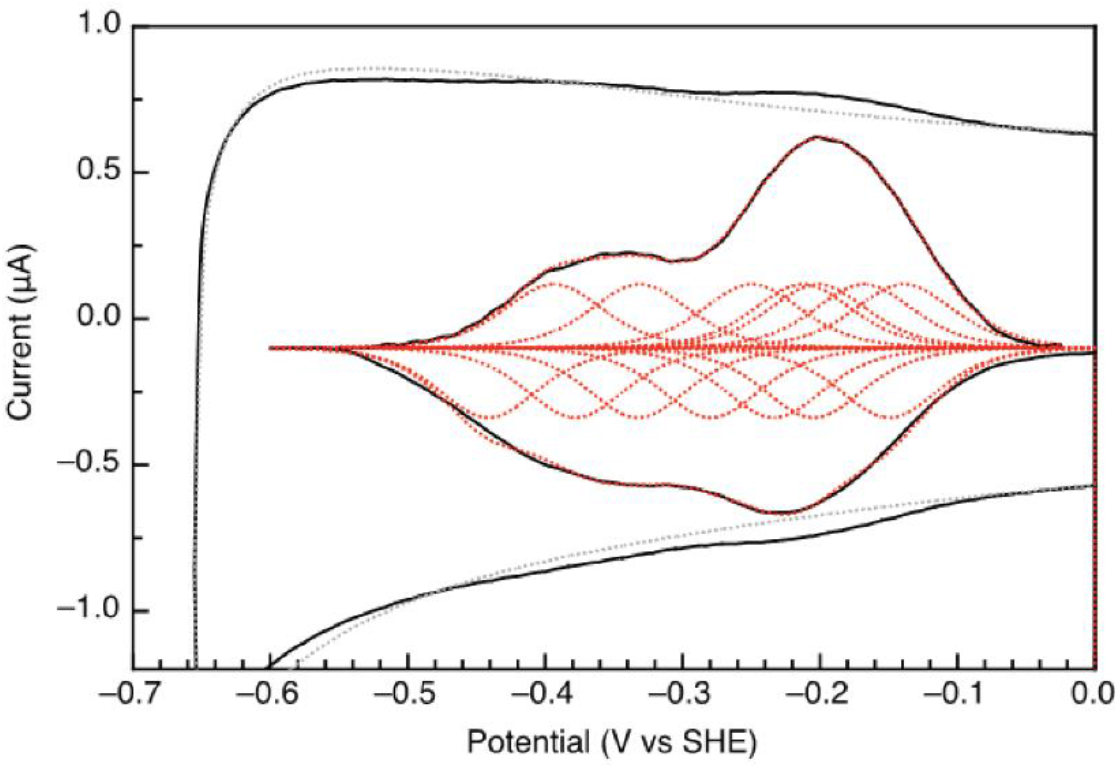
Electrochemical analysis of MmcA. Non-turnover MmcA voltammogram (black solid). Gray dotted line shows the ITO electrode baseline. The background-subtracted non-turnover MmcA voltammogram (black solid) with fitting (red dotted) are shown in the middle of the graph. Cyclic voltammetry (CV) recorded at pH 7.4, 10°C, and a scan rate of 20 mV/s.

**Supplementary Figure 13:**
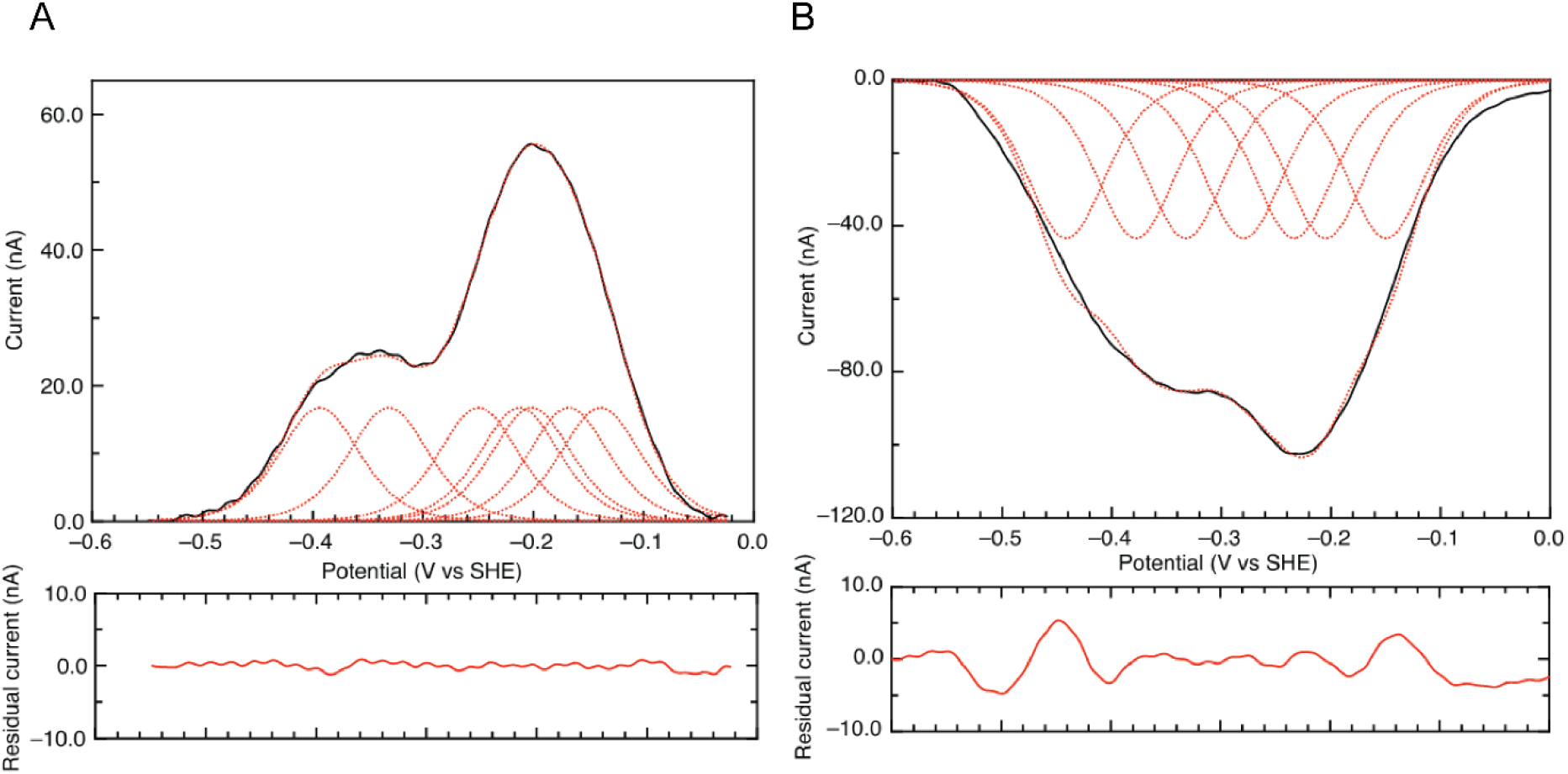
Electrochemical analysis of MmcA. Residuals for fits of the **(A)** oxidative and **(B)** reductive scans of MmcA. Cyclic voltammetry (CV) was recorded at pH 7.4, 10°C, and 20 mV/s.

**Supplementary Figure 14:**
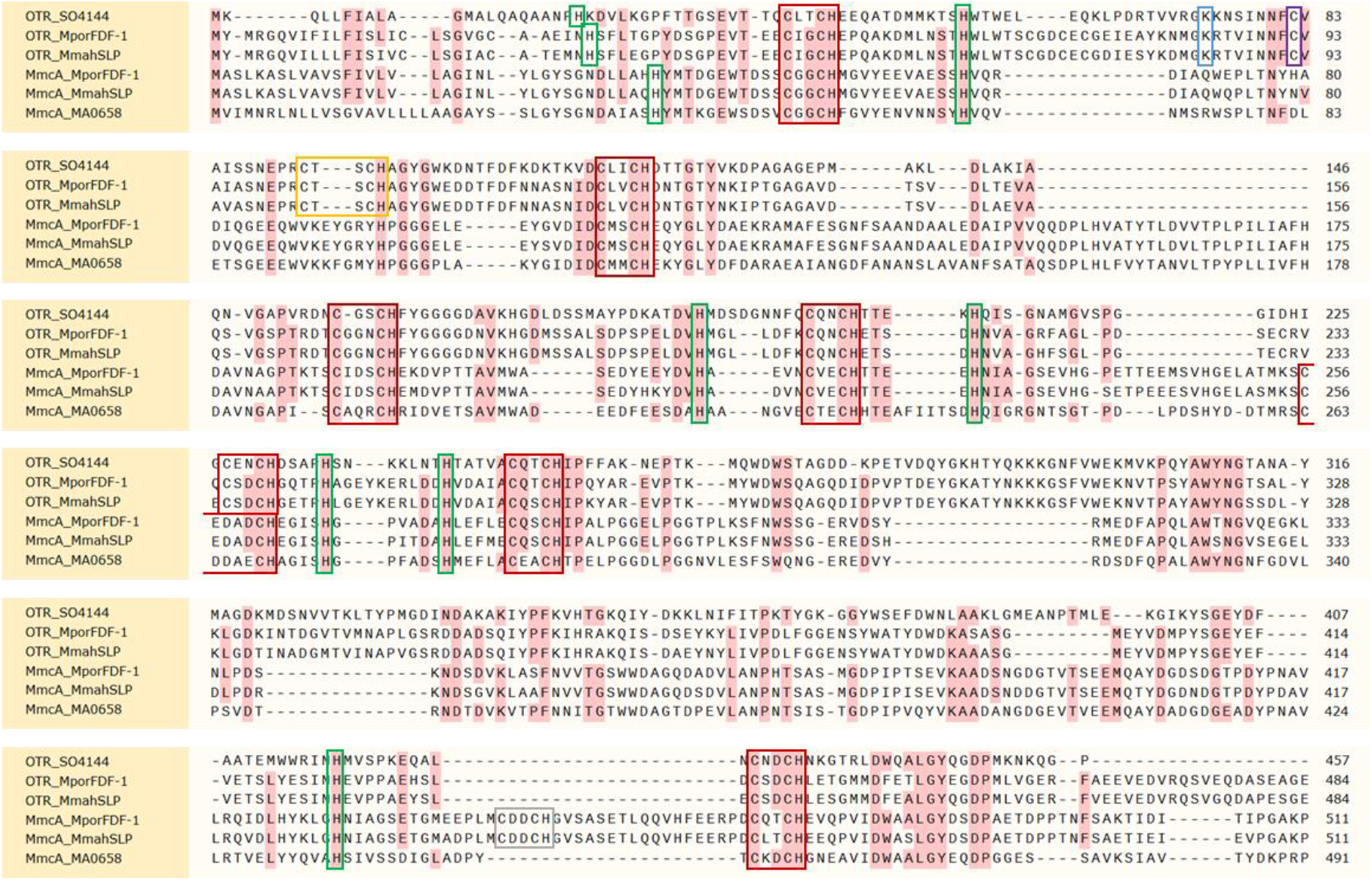
Protein alignment of MmcA and octaheme tetrathionate reductase (OTR) homologs. Alignment of octaheme tetrathionate reductase (OTR) from *Shewanella oneidensis* MR-1 (OTR_SO4144), MmcA from *M. acetivorans* (MmcA_MA0658), OTR and MmcA from *Methanohalophilus mahii* strain SLP (MmahSLP) as well as OTR and MmcA from *Methanohalophilus portucalensis* strain FDF-1 (MporFDF-1). The conserved features of the OTR catalytic site i.e., the second heme-binding motif (orange box), the lysine ligand (blue box) and the cysteine residue (purple box) are shown. Seven bis-His ligated heme-binding motifs (red box) and their distal histidine ligands (green box) in OTR and MmcA are also shown. An additional heme binding motif in MmcA from MmahSLP and MporFDF-1 (gray box) is highlighted.

**Supplementary Figure 15:**
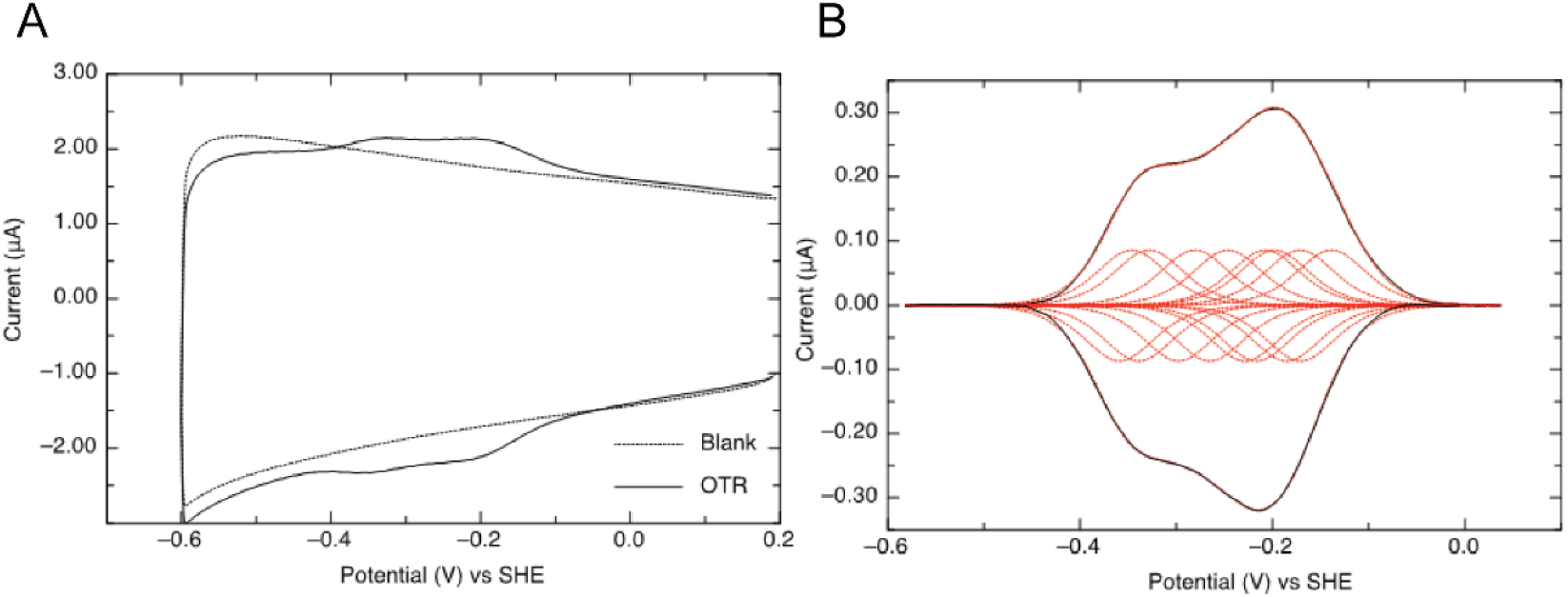
Electrochemical analysis of octaheme tetrathionate reductase (OTR) from *Shewanella oneidensis*. (**A**) Non-turnover OTR voltammogram (black solid). Dotted line shows the ITO electrode baseline. (**B**) Background-subtracted voltammogram with fitting of eight reversible redox couples (red dotted). Cyclic voltammetry (CV) recorded at pH 7.4, 10°C, and a scan rate of 20 mV/s.

**Supplementary Figure 16:**
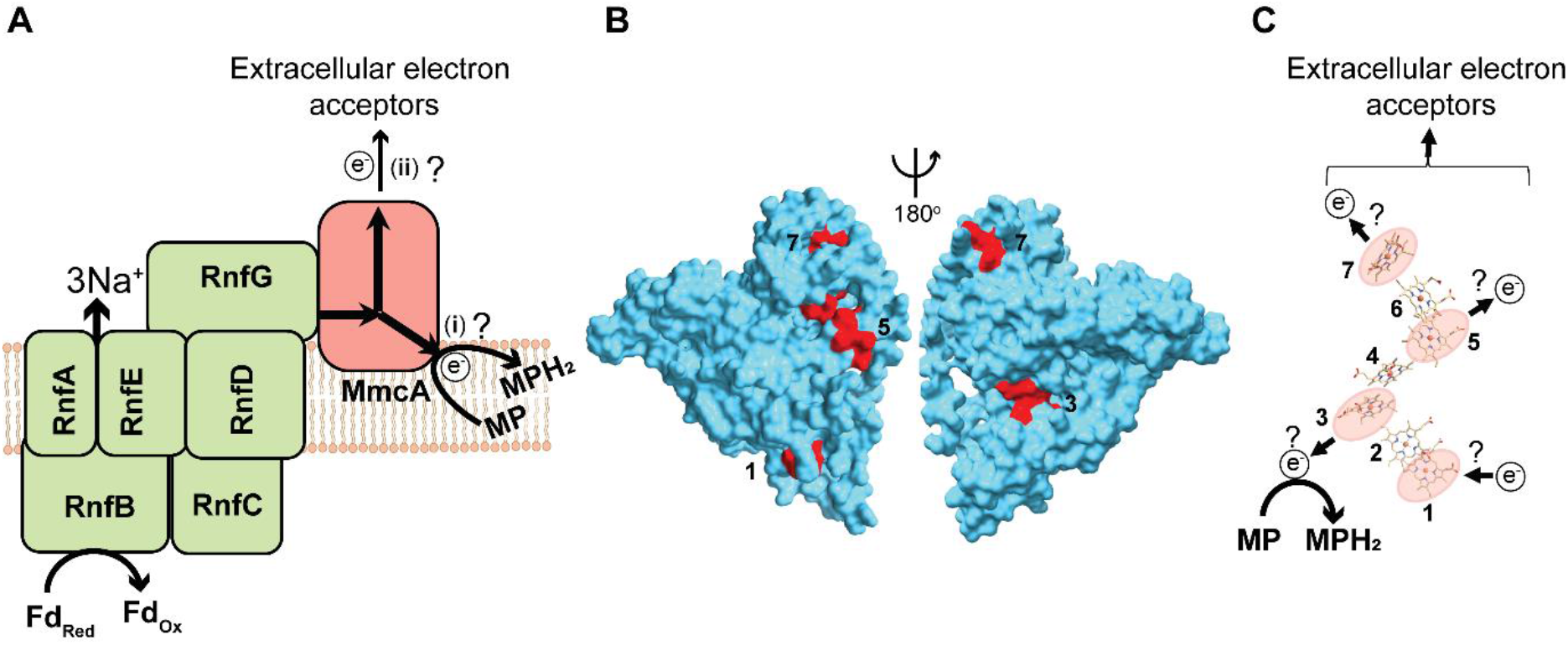
Proposed routes for electron transfer mediated by the heme groups in MmcA. (**A**) Schematic of the MmcA-Rnf complex where MmcA mediates the final step in the transfer of electrons from the complex to (i) the membrane-bound electron carrier methanophenazine (MP) and (ii) extracellular electron acceptors. (**B**) Surface of the AlphaFold predicted model of MmcA showing surface exposed heme-binding motifs i.e., 1, 3, 5 and 7 (red). (**C**) A plausible model of electron flow through MmcA. Electrons from Rnf complex could enter MmcA at a surface-exposed heme (such as heme #1) and be transferred to MP through another surface-exposed heme (such as heme #3). In the presence of extracellular electron acceptors, MmcA could simultaneously transfer electrons to extracellular electron acceptors through another surface-exposed heme (such as hemes #5 or #7). Surface-exposed heme groups (1, 3, 5 and 7) from panel B are highlighted in red circles. Although the model shows MmcA in a particular orientation, alternate orientations are equally consistent with our model.

**Supplementary Table 1:** Free energy change associated with methanogenesis and dissimilatory metal reduction (DSMR)

| Reaction | Electron donor | Electron acceptor | $\Delta G^{0'}$ per mol substrate |
| --- | --- | --- | --- |
| $\text{CH}_3\text{OH} \rightarrow \frac{3}{4} \text{CH}_4 + \frac{1}{4} \text{HCO}_3^- + \frac{1}{4} \text{H}^+$ | $\text{CH}_3\text{OH}$ | $\text{CH}_3\text{OH}$ | −78.75 kJ/mol |
| $\text{CH}_3\text{OH} + 6\text{Fe}^{3+} \rightarrow \text{HCO}_3^- + 6\text{Fe}^{2+} + 7\text{H}^+$ | $\text{CH}_3\text{OH}$ | $\text{Fe}^{3+}$ | −665.25 kJ/mol |
| $\text{CH}_3\text{COOH} \rightarrow \text{CH}_4 + \text{HCO}_3^- + \text{H}^+$ | $\text{CH}_3\text{COOH}$ | $\text{CH}_3\text{COOH}$ | −36 kJ/mol |
| $\text{CH}_3\text{COOH} + 8\text{Fe}^{3+} \rightarrow 2\text{HCO}_3^- + 8\text{Fe}^{2+} + 8\text{H}^+$ | $\text{CH}_3\text{COOH}$ | $\text{Fe}^{3+}$ | −810.44 kJ/mol |

$\Delta G^{0'}$ values were calculated from Decker et al., 1970.

**Supplementary Table 2:** List of *Methanosarcina acetivorans* strains used in this study

| Strains | Genotype | Source |
| --- | --- | --- |
| WWM60 | $\Delta hpt::PmcrB-tetR$ | (Guss et al., 2008) |
| DDN009 | $\Delta hpt::PmcrB-tetR, \Delta mmcA$ | (Gupta et al., 2022) |
| DDN016 | DDN009/ pDN409<br>[*PmcrB(tetO4)-mmcA- C- TAP tag*] | (Gupta et al., 2022) |
| DDN037 | DDN009/pDPG010<br>[[*PmcrB(tetO4)-uidA*] | (Gupta et al., 2022) |
| DDN038 | WWM60/pDPG010<br>[*PmcrB(tetO4)-uidA*] | (Gupta et al., 2022) |
| DDN039 | WWM60/pDN409<br>[*PmcrB(tetO4)-mmcA-C- TAP tag*] | (Gupta et al., 2022) |

## Notes

### Competing Interest Statement

The authors have declared no competing interest.

